# Olfaction regulates organismal proteostasis and longevity via microRNA-dependent signaling

**DOI:** 10.1101/420687

**Authors:** Fabian Finger, Franziska Ottens, Alexander Springhorn, Tanja Drexel, Lucie Proksch, Sophia Metz, Luisa Cochella, Thorsten Hoppe

## Introductory Paragraph

The maintenance of proteostasis is crucial for any organism to survive and reproduce in an ever-changing environment, but its efficiency declines with age^1,2^. Posttranscriptional regulators such as microRNAs control protein translation of target mRNAs with major consequences for development, physiology, and longevity^3,4^. However, the precise function of lifespan-determining microRNAs remains poorly understood. Here we show that the microRNA *mir-71* controls organismal proteostasis and aging in *Caenorhabditis elegans* by regulating its conserved target *tir-1* in AWC olfactory neurons. We screened a collection of microRNAs that control aging^4^ to identify regulators of organismal proteostasis and discovered that the lifespan promoting *mir-71* affects ubiquitin-dependent protein turnover, particularly in the intestine. We show that *mir-71* directly inhibits the toll receptor domain protein TIR-1 in AWC olfactory neurons. Neuronal signaling is required for *mir-71*/*tir-1*-dependent and diet-dependent regulation of organismal proteostasis. Disruption of *mir-71*/*tir-1* or loss of AWC olfactory neurons eliminates the influence of food source on proteostasis. *Mir-71*-mediated regulation of TIR-1 controls chemotactic behavior and is regulated by odor. Our findings support a model whereby odor promotes *mir-71*-mediated inhibition of TIR-1 in AWC neurons to stimulate organismal protein turnover. Thus, odor perception influences cell-type specific miRNA-target interaction to regulate organismal proteostasis and longevity. We anticipate that the proposed mechanism of food perception will stimulate further research on neuroendocrine brain-to-gut communication and may open the possibility for therapeutic interventions to improve proteostasis and organismal health via the sense of smell, with potential implication for obesity, diabetes and aging.

MicroRNA biogenesis is associated with improved stress tolerance and longevity in *C. elegans*^5^, and specific microRNAs have been linked to stress resistance and aging^4,6,7^. Changes in stress resistance and longevity are often associated with alterations in protein homeostasis (proteostasis)^1,2^. To identify microRNAs that control longevity via effects on organismal proteostasis, we used established *in vivo* assays to monitor ubiquitin-mediated turnover of fluorescently labeled model substrates in *C. elegans*^8,9^. The UbV-GFP protein is a ubiquitously expressed, ubiquitin fusion degradation (UFD) substrate. In wild-type worms, poly-ubiquitylation of the N-terminal ubiquitin moiety triggers degradation of UbV-GFP by the 26S proteasome. Disruption of ubiquitylation or of proteasomal turnover results in UbV-GFP stabilization in transgenic worms (Fig. 1a)^8^. We also monitored endoplasmic reticulum (ER)-associated protein degradation (ERAD) by following the turnover of an unstable form of the cathepsin L-like cysteine protease CPL-1 (CPL-1*-YFP) expressed in intestinal cells^9^. CPL-1*-YFP is normally retro-translocated out of the ER lumen for ubiquitin-mediated proteasomal degradation in the cytosol; loss of the ERAD-associated E3 ubiquitin-protein ligase SEL-1 triggers substrate stabilization (Supplementary Fig. 1f, 3d)^10^.

**Figure 1.**
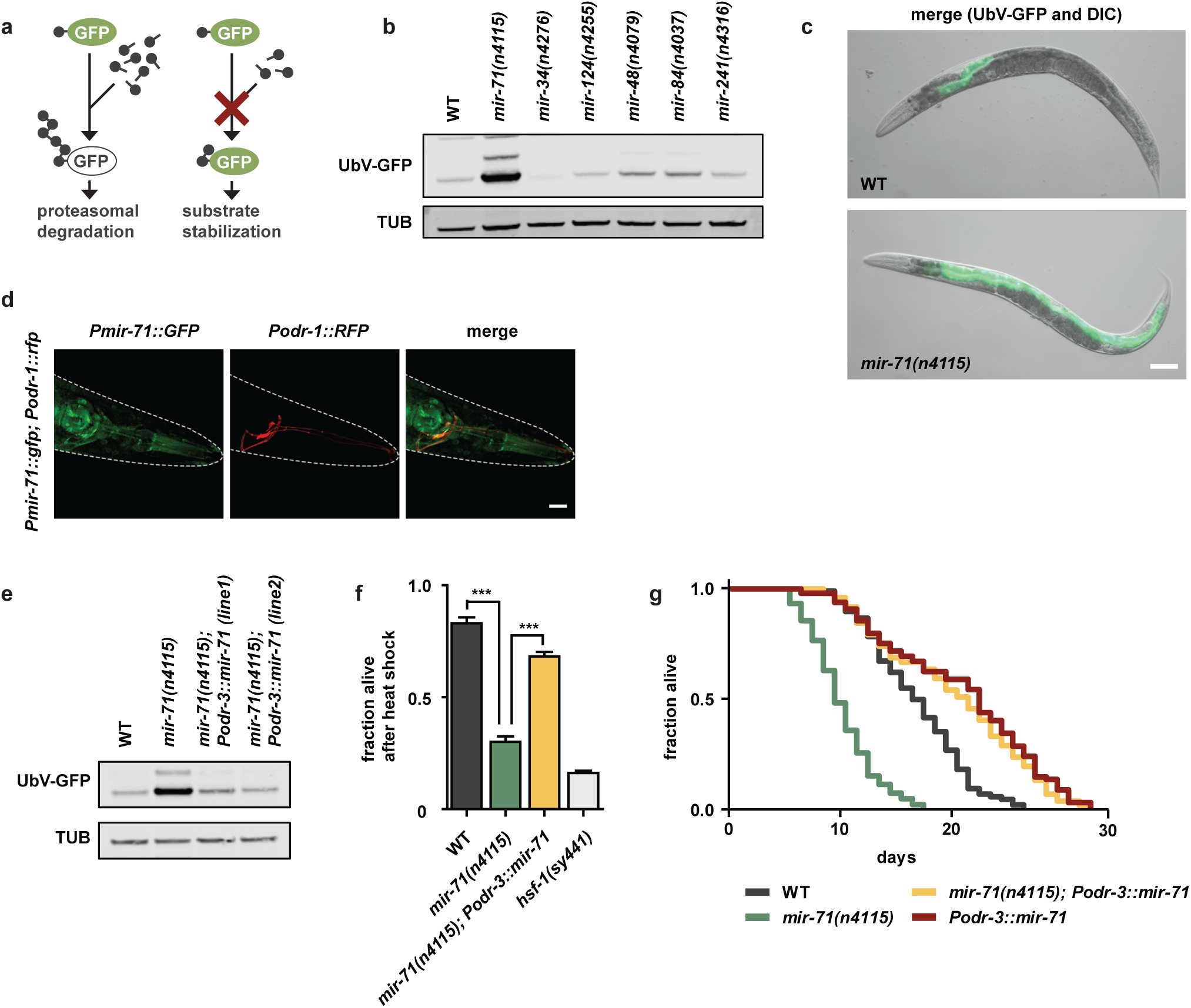
| Expression of *mir-71* in olfactory neurons supports proteostasis and longevity. **a**, The ubiquitin fusion degradation (UFD) model substrate for monitoring ubiquitin-dependent degradation. **b**, The *mir-71(n4115)* deletion allele exhibits stabilization of the UFD substrate. Detection of the GFP signals via western blot showing UbV-GFP and tubulin (TUB) level. **c**, The UFD substrate accumulates mainly in the intestine upon *mir-71(n4115)* deletion. Representative fluorescent images of day 1 adult worms with indicated genotypes. Scale bar: 250 μm. **d**, *mir-71* is expressed in olfactory (AWC) neurons. Confocal microscopy images showing localization of *Pmir-71∷GFP* in green and AWC neurons in red. Scale bar: 15 µm. **e**, AWC-specific rescue of the *mir-71(n4115)* deletion mutant restores protein degradation. Western blot from day 1 adult worm lysates with indicated genotypes showing UbV-GFP and tubulin (TUB) levels. **f**, AWC-specific expression of *mir-71* increases survival upon heat stress. The *hsf-1(sy441)* mutant served as control. Data represent mean values ± SEM generated from n=3 independent experiments using at least 50 worms; ***p < 0.001, ns=not significant. **g**, AWC-specific expression of *mir-71* extends lifespan. For statistics details see Supplementary Table 1.

Of the microRNA loss-of-function mutants we tested, we found that *mir-71(n4115)* showed a substantial increase in both UbV-GFP and CPL-1*-YFP levels, particularly within the intestine, relative to wild-type worms (Fig. 1b, c, Supplementary Fig. 1a, g, h). Importantly, the levels of ubiquitylation-resistant ^K^29^/^48^R^UbV-GFP and GFP were unaltered in *mir-71(n4115)* (Supplementary Fig. 1b)^8^, and wild-type and *mir-71(n4115)* animals showed comparable levels of *UbV-GFP* mRNA (Supplementary Fig. 1c). Overexpression of the proteasomal subunit RPN-6.1, which triggers degradation of ubiquitylated proteins^11^, suppressed the stabilization of UbV-GFP in *mir-71(n4115)* (Supplementary Fig. 1d). In contrast, loss of the E3 ligase HECD-1, which acts upstream of the 26S proteasome in substrate ubiquitylation, cannot be compensated by elevated RPN-6.1 level (Supplementary Fig. 1d)^8^. RNAi-mediated knockdown of *hecd-1,* as well as *rpn-8,* led to an additive defect when combined with *mir-71(n4115)* (Supplementary Fig. 1e). These data suggest that *mir-71* regulates ubiquitin-dependent protein degradation via the 26S proteasome.

Consistent with ERAD defects, *mir-71(n4115)* worms are sensitive to ER stress induced by tunicamycin (TM), which blocks N-linked glycosylation of ER proteins^12^. Interestingly, the previously reported lifespan reduction of *mir-71(n4115)*^4^ was further decreased by TM treatment (Supplementary Fig. 1i). To further characterize the role of *mir-71* in intestinal proteostasis, we performed tissue-specific rescue experiments^13^. Ubiquitous and pan-neuronal, but not hypodermal, expression of *mir-71* in *mir-71(n4115)* rescued defective turnover of UbV-GFP (Supplementary Fig. 2a, b). These data suggest that loss of *mir-71* expression in neurons stabilizes UbV-GFP in intestinal cells. Previous work found that *mir-71* is expressed in olfactory neurons^14^. To identify the type of neuron required for *mir-71*-dependent regulation of proteostasis, we genetically abolished the specification of AWA, AWB/ADF, AWC, or PQR/PHA/PHB olfactory neurons by depleting the transcription factors required for their cell-fate determination, ODR-7, LIM-4, CEH-36, and CEH-14, respectively^15^. Abrogation of either AWB/*lim-4* or AWC/*ceh-36* development partially suppressed UbV-GFP accumulation in *mir-71(n4115)* (Supplementary Fig. 2c), suggesting that these olfactory neurons are required for *mir-71*-mediated regulation of intestinal homeostasis. Intriguingly, the AWCs belong to a class of ciliated olfactory neurons previously implicated in the regulation of longevity^16^. This led us to hypothesize a prominent role for AWC neurons in organismal proteostasis via *mir-71*.

To test this hypothesis, we first confirmed that *mir-71* is expressed in AWC neurons. Indeed, we observed colocalization of *Pmir-71∷GFP* and the AWC-specific marker *Podr-1∷RFP* (Fig. 1d)^14^. Further, *Podr-1∷RFP* expression revealed comparable AWC neuronal integrity in *mir-71(n4115)* and wild-type worms (Supplementary Fig. 2d). Intriguingly, exclusive expression of *mir-71* in AWC (*Podr-3*) but not AWB neurons (*Pstr-1*), fully restored the ability of *mir-71(n4115)* animals to degrade UbV-GFP (Fig. 1e, Supplementary Fig. 2e, f)^17,18^. AWC-specific expression of *mir-71* in *mir-71(n4115)* alleviated mortality induced by the proteasome inhibitor bortezomib (BTZ) (Supplementary Fig. 2g), as well as heat sensitivity and reduced lifespan (Fig. 1f, g). Further, *Podr-3∷mir-71* expression prolonged lifespan of wild-type worms (Fig. 1g), suggesting that *mir-71* improves organismal physiology. Thus, *mir-71* expression in AWC olfactory neurons is necessary and sufficient to coordinate organismal proteostasis, particularly in the intestine.

MicroRNAs regulate gene expression via complementary base pairing with target mRNAs. To identify potential targets of *mir-71* important for proteostasis regulation, we compiled a list of genes that were differentially expressed in olfactory neurons (i) upon proteasomal inhibition of wild-type worms, and (ii) in the *mir-71(n4115)* deletion mutant, relative to wild-type (Supplementary Fig. 3a, Supplementary Table 2). We further reduced the list to AWC-specific mRNAs regulated in response to proteotoxic stress, based on RNA sequencing data and bioinformatic microRNA target prediction (Fig. 2a, Supplementary Table 3) (TargetScanWorm Release 6.2;^19^).

**Figure 2.**
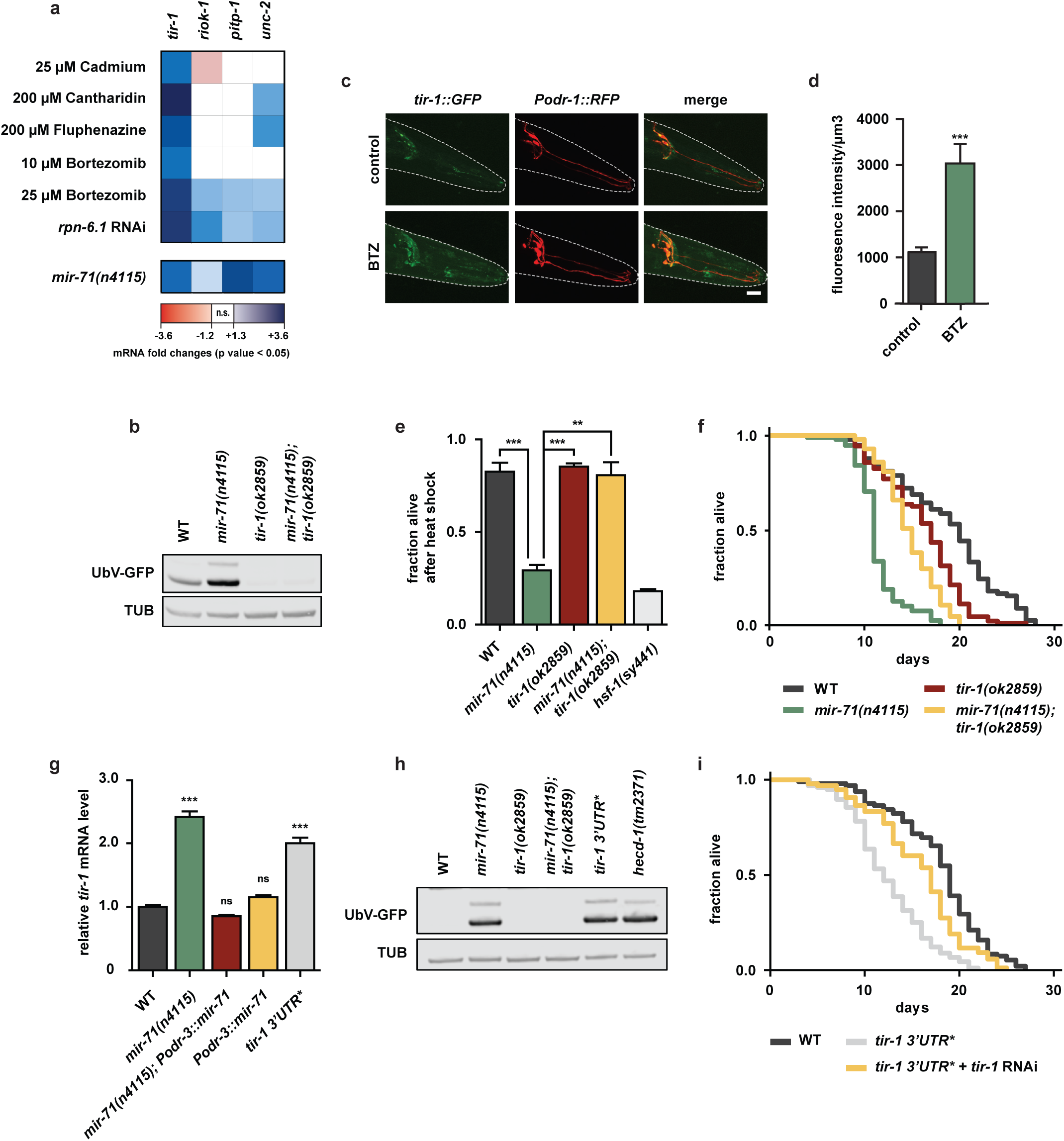
| Negative regulation of *tir-1* by *mir-71* important for proteostasis and longevity. **a**, mRNA level of predicted *mir-71* targets (TargetScanWorm Release 6.2). Color-coded transcriptional responses of mRNAs expressed in AWC neurons under multiple proteotoxic stress conditions or in *mir-71(n4115)* deletion mutant. Expression levels are relative to untreated wild-type (WT) worms and were determined via RNA sequencing (upper panel) or microarray (lower panel), n.s.=not significant. **b**, **h**, Representative western blots from day 1 adult worms of indicated genotypes show UbV-GFP and tubulin (TUB) level. **b**, The *tir-1(ok2859)* deletion mutant suppresses *mir-71(n4115)*-induced proteolytic defects. **c**, Proteotoxic stress increases TIR-1 levels in AWC neurons. Confocal microscopy images showing *tir-1∷GFP* in green and AWC neurons in red. Scale bar: 15 µm. **d**, Quantification of TIR-1∷GFP fluorescence intensity shown in **c**. DMSO was used as a solvent control but did not show a significant difference compared to the control condition (1322.24 ± 96.05, data not shown). Data represent mean values ± SEM generated from imaging n=12 individual animals; ***p < 0.001. **e**, Heat stress sensitivity of *mir-71(n4115)* mutant is suppressed by *tir-1(ok2859)* deletion. The *hsf-1(sy441)* mutant served as control. Data show mean values ± SEM obtained from n=3 independent experiments using at least 50 worms; **p < 0.01, ***p < 0.001. **f**, *tir-1(ok2859)* deletion extends the lifespan of *mir-71(n4115)* worms. For statistics details see Supplementary Table 1. **g**, *mir-71* negatively regulates *tir-1* transcript level. Relative *tir-1* mRNA levels measured by qRT-PCR in day 1 adult worms. Data shown as mean values ± SEM generated from n=3 technical replicates; values are normalized to wild-type (WT); ***p < 0.001, ns=not significant. **h**, AWC-specific expression of *mir-71* does not suppress proteostasis defects caused by the *tir-1 3’UTR\** mutation. **i**, The *tir-1 3’UTR\** mutation shortens lifespan. For statistics details see Supplementary Table 1.

We identified *tir-1*, *riok-1*, *pitp-1*, and *unc-2* as potential targets of *mir-71*. In particular, *tir-1* mRNA was strongly elevated in response to proteotoxic reagents and contains three predicted *mir-71* binding sites within the 3’UTR (Fig. 2a, Supplementary Fig. 3a, Supplementary Table 2, 3). To test whether up-regulation of the identified genes might account for the defects in proteostasis of *mir-71*-deficient animals, we depleted the candidate mRNAs by RNAi. Only *tir-1* depletion restored UbV-GFP levels in *mir-71(n4115)* mutants (Supplementary Fig. 3b). Further, degradation of both the UFD and ERAD substrates was restored in *mir-71(n4115)*; *tir-1(ok2859)* double mutants (Fig. 2b, Supplementary Fig. 3c, d). Finally, ubiquitylation-resistant ^K^29^/^48^R^UbV-GFP or GFP proteins, as well as *UbV-GFP* mRNA levels, were unaltered in *tir-1(ok2859)* (Supplementary Fig. 3e, f), revealing that the decrease in substrate protein levels did not result from transcriptional or translational changes.

TIR-1 is a toll receptor domain adapter protein that is important for innate immune responses and neuronal development^14,20,21^. To determine where *tir-1* is expressed, we used CRISPR-Cas9-based gene editing to generate a *tir-1∷GFP* reporter. We observed TIR-1∷GFP expression in AWC olfactory neurons^14,20^. Further, proteotoxic stress via BTZ treatment increased the expression of TIR-1∷GFP as well as *tir-1* mRNA levels (Fig. 2a, c, d). Importantly, loss of *tir-1* suppressed the heat stress sensitivity and the shortened lifespan of *mir-71(n4115)* mutants (Fig. 2e, f). We also evaluated a gain-of-function mutant *tir-1(yz68)*, which mediates constitutive activation of TIR-1^22^. In contrast to loss of TIR-1, the *tir-1(yz68)* mutant caused a slight increase in UbV-GFP levels, which further increased in the context of *mir-71(n4115)* (Supplementary Fig. 3g, h). These findings suggest that *mir-71* negatively regulates *tir-1*.

To test whether *tir-1* mRNA is a direct target of *mir-71,* we used a CRISPR/Cas9 approach to disrupt all three predicted *mir-71* binding sites in an endogenous allele of *tir-1* in wild-type worms (*tir-1 3’UTR\**) (Supplementary Fig. 4a). Indeed, *tir-1* mRNA levels increased in *tir-1 3’UTR\** worms relative to wild-type (Fig. 2g, grey bar), indicating a direct repressive effect of the microRNA. Further, the *mir-71(n4115)* deletion mutant showed increased levels of *tir-1* mRNA (Fig. 2g, green bar), in line with our microarray analysis (Fig. 2a). Expression of *mir-71* in AWCs restored *tir-1* mRNA levels back to wild-type in *mir-71(n4115)*, demonstrating that the *tir-1* de-regulation observed in *mir-71* mutant animals occurs primarily in AWC neurons (Fig. 2g, green versus red bar).

Importantly, animals carrying the *tir-1 3’UTR\** allele accumulated both UFD and ERAD substrates, fully phenocopying *mir-71(n4115)*-related defects in protein degradation (Fig. 2h, Supplementary Fig. 4b-d). RNAi-mediated depletion of *tir-1* or overexpression of RPN-6.1 suppressed the protein turnover defect in *tir-1 3’UTR\** animals (Supplementary Fig. 4e, f). Thus, the substrate stabilization caused by *tir-1 3’UTR\** depends on increased *tir-1* levels and results from ubiquitin proteasome system (UPS) dysfunction. Overexpression of *mir-71* in AWC neurons did not re-establish UPS function in *tir-1 3’UTR\** animals, highlighting the unique importance of *tir-1* mRNA regulation for proteostasis (Supplementary Fig. 4g). Similar to loss of *mir-71*, the *tir-1 3’UTR\** mutation also reduced lifespan, which was rescued by RNAi-mediated depletion of *tir-1* (Fig. 2i). These data suggest that *mir-71/tir-1* represents a one-to-one microRNA/mRNA interaction that regulates proteome stability and longevity.

We previously showed that feeding worms with different bacterial food sources modifies protein degradation in the intestine^8^. As expected, UbV-GFP was efficiently degraded in wild-type worms grown on standard food, *Escherichia coli* (*E. coli*) strain OP50, but accumulated in worms fed *E. coli* C600, HT115, or HB101 bacteria (Fig. 3a). In contrast, *mir-71(n4115)* mutants did not display food-related changes in ubiquitin-dependent proteolysis, and this was restored by AWC-specific *mir-71* expression (Fig. 3a, b, Supplementary Fig. 5a).

**Figure 3.**
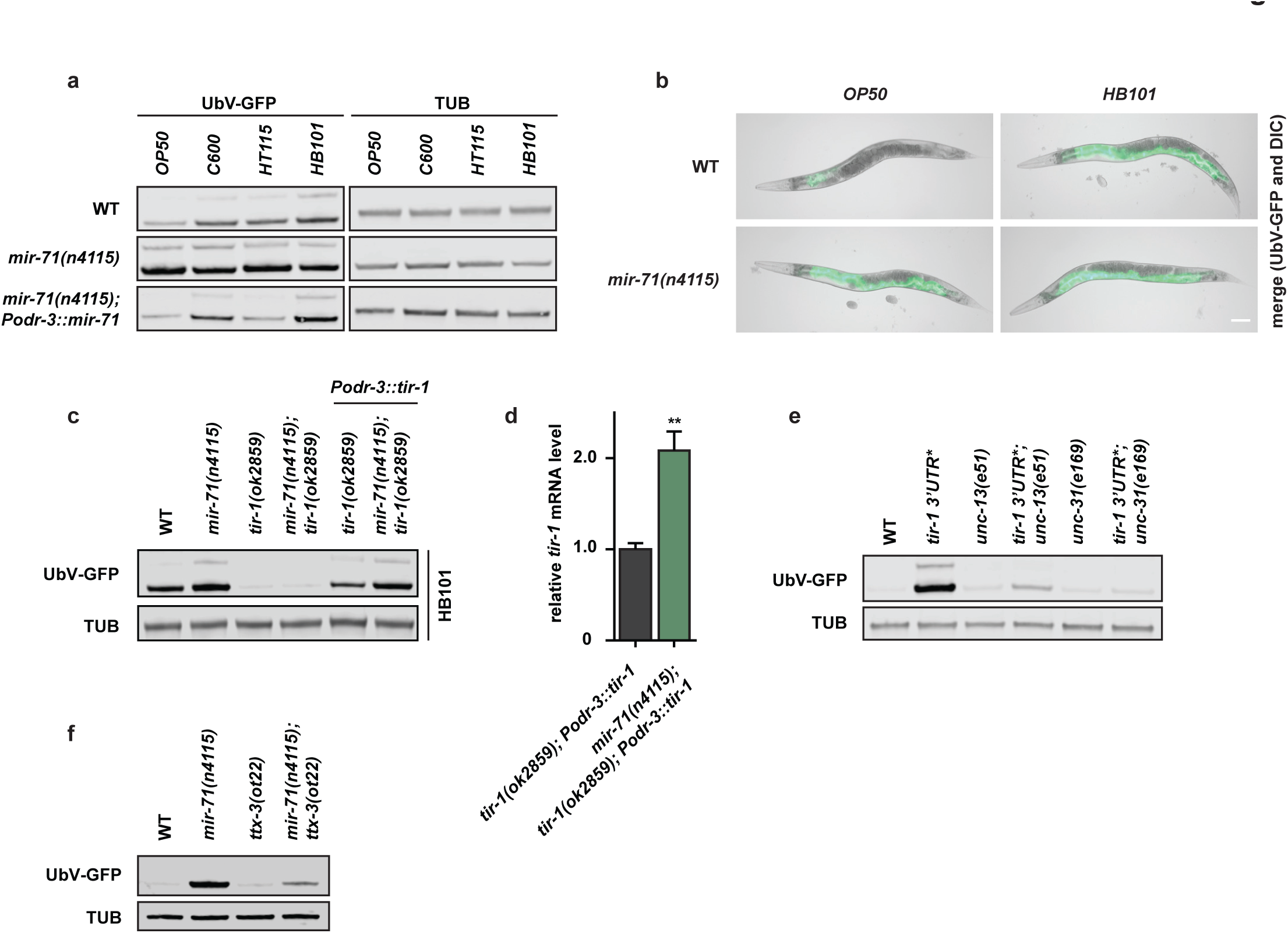
| Food-dependent coordination of proteostasis is triggered by *mir-71/tir-1* dynamics in AWC neurons. **a**, **c**, **e**, **f,** Representative western blots of worm lysates with the indicated genotypes show UbV-GFP and tubulin (TUB) level. **a**, AWC-specific *mir-71* expression is important for food perception and proteostasis. Animals were grown on OP50, C600, HT115, or HB101 bacteria prior to worm lysis. **b**, Representative fluorescent images of day 1 adult worms grown on OP50 or HB101 with indicated genotypes. Scale bar: 250 μm. **c**, AWC-specific *tir-1* expression rescues food-dependent changes in protein degradation. **d**, *mir-71* negatively regulates *tir-1* transcript level in AWC neurons. Relative *tir-1* mRNA levels measured by quantitative real-time PCR (qRT-PCR) in day 1 adult worms. Data shown as mean values ± SEM generated from n=3 technical replicates; values are normalized to *tir-1(ok2859); Podr-3∷tir-1*; **p < 0.01, ns=not significant. **e**, Neuronal signaling is required for changes in UPS activity caused by the *tir-1 3’UTR\** mutation. **f**, Reduced AIY interneuron activity caused by *ttx-3(ot22)* suppresses protein degradation defects of *mir-71* deletion.

Similar to *mir-71(n4115)*, *tir-1 3’UTR\** animals exhibited high levels of UbV-GFP, irrespective of different bacterial strains (Supplementary Fig. 5b). In contrast, *tir-1(ok2859)* loss-of-function mutants did not accumulate UbV-GFP when grown on HB101, which was reversed upon AWC-specific *tir-1* expression (Fig. 3c, d, Supplementary Fig. 5b, c). We found that *UbV-GFP* and *tir-1* mRNA levels were higher when *Podr-3∷tir-1* was expressed in *mir-71(n4115)* mutants compared to wild-type worms, consistent with loss of miRNA-mediated *tir-1* suppression in *mir-71(n4115)* (Fig. 3c, d, Supplementary Fig. 5c). These data suggest that food-related changes in proteostasis are mediated at least in part by *mir-71*-dependent, cell type-specific regulation of *tir-1* in AWC olfactory neurons.

Cell-nonautonomous, neuroendocrine signaling pathways have been shown to trigger organismal regulation of proteotoxic stress and quality control^23–25^. Since UbV-GFP accumulated mainly in the intestine upon exposure to different bacteria or *mir-71* deletion, we tested whether AWC-specific *tir-1* regulation is communicated to peripheral tissues by neuronal signaling. We used an UNC-13 mutant to evaluate the consequence of blocking neurotransmitter release from small clear vesicles (SCV), and an UNC-31 mutant to block neuropeptide release from dense core vesicles (DCV)^26,27^. We found that both *unc-13(e51)* and *unc-31(e168*) suppressed the substrate stabilization caused by feeding HB101 bacteria (Supplementary Fig. 5a), by *mir-71* loss-of-function (Supplementary Fig. 5d), and by *tir-1 3’UTR\** (Fig. 3e), suggesting that neuronal signal transduction is important for AWC-dependent coordination of proteostasis in peripheral tissues.

To determine the signaling molecules that trigger the cell-nonautonomous communication from the olfactory neurons to the intestine, we evaluated changes in the expression of neuropeptide genes in *mir-71(n4115)* compared to wild-type worms (Supplementary Fig. 5e and Supplementary Table 4)^28^. We observed high upregulation of NLP-9 and NLP-14 in *mir-71(n4115)* worms (Supplementary Fig. 5e). Further, RNAi-mediated depletion of NLP-9 and NLP-14 partially or fully suppressed *mir-71(n4115)*-dependent degradation defects, respectively (Supplementary Fig. 5f). Thus, these data suggest that AWC neurons regulate intestinal homeostasis via secretion of the neuropeptides NLP-9 and NLP-14.

AWC neurons communicate with AIB and AIY interneurons, among others, to initiate food- and odor-derived behavior^29^. To validate the role of *mir-71* for communication via the olfactory system, we analyzed a loss-of-function mutant for *ttx-3*, which encodes a LIM homeodomain protein necessary for AIY function^30^. Indeed, the *mir-71(n4115)*-dependent defects in protein degradation were suppressed in the *ttx-3(ot22)* mutant background, supporting the idea that signal transduction via AIY interneurons connects food-derived olfactory inputs and proteostasis (Fig. 3f).

To define the cause of food-related changes in proteostasis, we monitored degradation of UbV-GFP in wild-type worms exposed to mixtures of OP50 and HB101 bacteria. In contrast to HB101 alone, worms raised on HB101/OP50 mixtures (100:1, 10:1, and 1:1) showed significantly decreased substrate stabilization almost indistinguishable from worms exclusively exposed to OP50 (Fig. 4a). However, *mir-71(n4115)* and *tir-1 3’UTR\** worms did not distinguish the different mixtures and showed UbV-GFP substrate stabilization, which did not exceed effects with HB101 alone in all cases (Fig. 4a). The very low detection threshold for the proteostasis response to OP50 is consistent with *C. elegans’* ability to detect low concentrations of volatile chemicals^31^. To acutely silence AWC activity specifically after development, we expressed a histamine-gated chloride channel (HisCl1) under control of the *ceh-36* promoter and treated worms with histamine (Fig. 4b)^32^. Histamine (HA) treatment of worms grown on OP50 or HB101 bacteria completely abolished the food-evoked differences in UbV-GFP stabilization (Fig. 4c). Strikingly, HA-treatment similarly suppressed degradation defects of *mir-71(n4115)* worms on both bacterial strains, which suggests a direct link between neuronal activity and *mir-71* function in food-dependent regulation of proteostasis (Fig. 4d).

**Figure 4.**
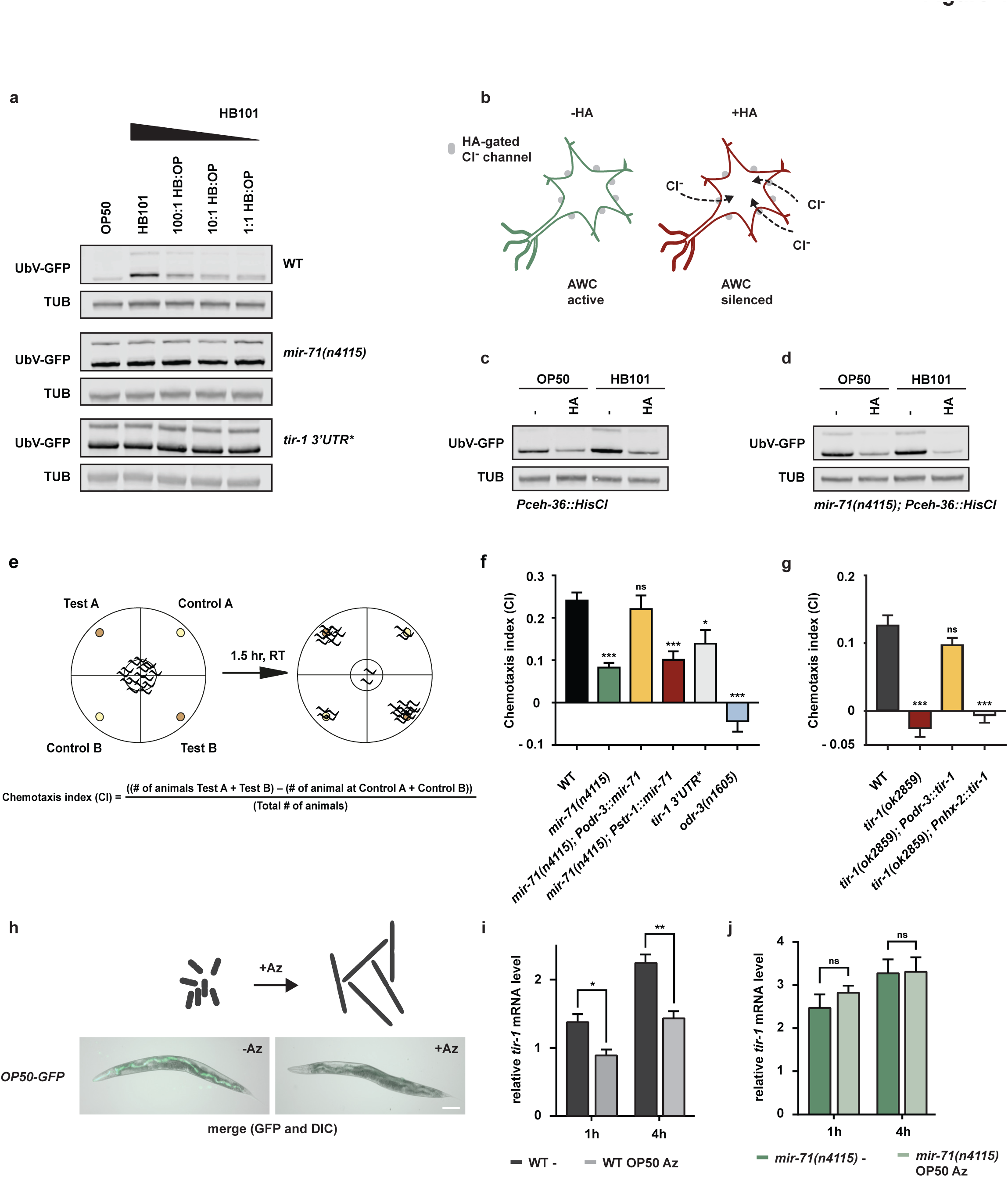
| Coordination of food perception and organismal proteostasis. **a**, **c**, **d**, Representative western blots showing UbV-GFP and tubulin (TUB). **a**, Low concentrations of OP50 alleviate HB101-dependent changes in proteostasis. Worm strains of indicated genotype were grown on depicted mixture of HB101 and OP50. **b**, AWC silencing by histamine-gated chloride channel expression (*Pceh-36∷HisCl*). **c-d**, Food-dependent proteostasis defects in WT (**c**) and *mir-71(n4115)* (**d**) are inhibited upon AWC silencing. Age-synchronized L4 larvae were grown on control (-) or histamine (HA) containing plates (10 mM HA) until day 1 of adulthood. **e**, Schematic overview of the chemotaxis assay: Plates are separated into two test (Test A and B) and two control (Control A and B) quadrants. A drop of bacterial culture was placed into the test quadrants. Worms were transferred to the center of the plate and after a 1.5 hr incubation at room temperature (RT) the number of animals in each quadrant was determined. Calculation of the chemotaxis index (CI) is shown below. **f**, **g**, AWC-specific expression of *tir-1* or *mir-71* is important for chemotaxis behavior towards bacterial food source (*E. coli* strain OP50). Chemotaxis assays were performed as depicted in Fig. 4e using worm strains of the indicated genotypes. Data presented as mean values ± SEM generated from n=9 independent experiments with at least 66 (**f**) or 80 animals (**g**); values are normalized to wild-type (WT); *p < 0.05, **p < 0.01***p < 0.001, ns=not significant. **h**, *E. coli* bacteria treated with Aztreonam (Az) show abnormal cell growth and cannot be ingested by *C. elegans.* For ingestion control *E.coli* OP50 expressing GFP (OP50-GFP) were either treated with Az or left untreated before seeding them to NGM plates. Worms were grown on the respective bacteria for 6 hr prior to imaging. Representative fluorescent images of day 1 adult worms. Scale bar: 250 μm. **i**, **j**, The smell of food affects *tir-1* mRNA level. Relative *tir-1* mRNA levels measured by qRT-PCR in day 1 adult worms. Age-synchronized worms were grown on OP50 bacteria until day 1 of adulthood and then transferred to plates without food (-) or Az-treated OP50 (OP50 Az) (see also Fig. 4h) for 1 h or 4 hr. Data shown as mean values ± SEM generated from n=3 technical replicates; values are relative to wild-type (WT) *tir-1* level on food (control condition, data not shown); statistics between no food (-) and OP50 Az; **p < 0.01, *p < 0.05, ns=not significant.

To discern the role of AWC neurons in smelling volatile odors, we used standard chemotaxis assays to test whether *tir-1* regulation is necessary for sensing a bacterial food source (Fig. 4e)^33^. Indeed, both *mir-71(n4115)* and *tir-1 3’UTR\** animals showed reduced food detection. The compromised chemosensation of *mir-71(n4115)* could be rescued by *mir-71* expression in AWC but not AWB neurons (Fig. 4f, Supplementary Table 5). Similarly, chemotaxis defects of *tir-1(ok2859)* deletion animals were rescued by AWC-specific *tir-1* but not intestinal expression (Fig. 4g, Supplementary Table 5), suggesting that cell type-specific *tir-1* regulation is required both for olfactory food perception and proteostasis. To test this hypothesis, we measured the level of *tir-1* mRNA in wild-type worms exposed to the smell of bacteria. We incubated OP50 bacteria with aztreonam (Az), an antibiotic that affects bacterial cell division, resulting in inedible bacterial filaments (Fig. 4h)^34^. We found that *tir-1* mRNA levels increased in wild-type worms transferred to a plate without food, and the smell of inedible bacteria suppressed *tir-1* elevation in this context (Fig. 4i). Interestingly, *mir-71(n4115)* mutants, which have already upregulated *tir-1* mRNA, did not further increase in *tir-1* mRNA levels upon transfer to a plate lacking food (Fig. 4j). These data suggest that the absence of an odor relieves *mir-71* mediated repression of *tir-1* in wild-type worms. Our findings support a direct link between olfactory food perception and *mir-71*-mediated inhibition of TIR-1 to control proteostasis and aging (Fig. 5).

**Figure 5.**
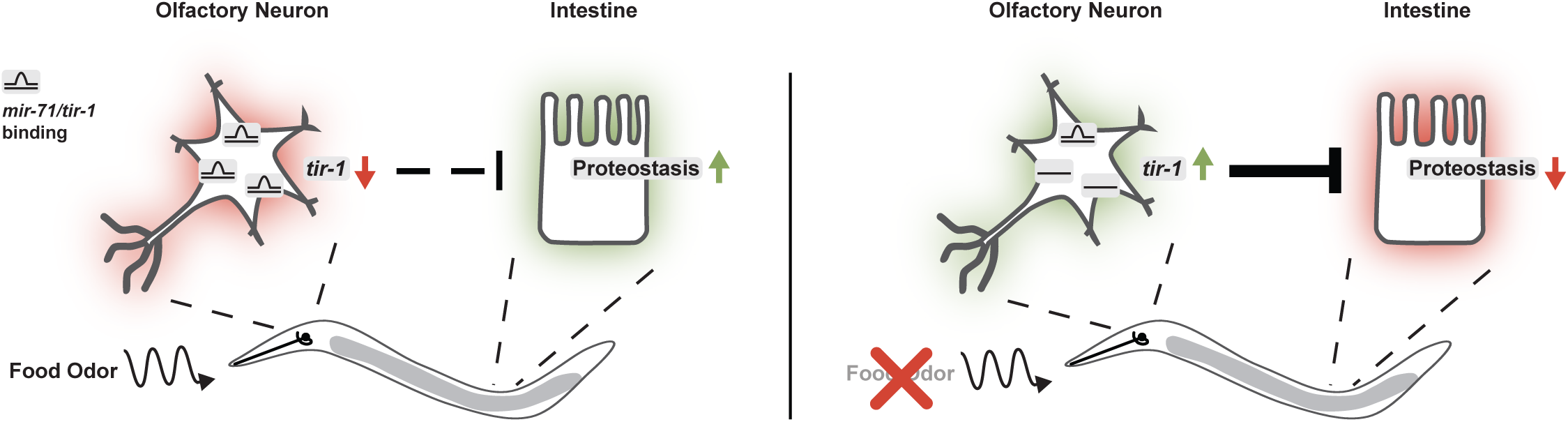
| Model. Odors related to bacterial food are sensed via ciliated AWC olfactory neurons. The olfactory stimulation of AWCs provokes cell-nonautonomous regulation of ubiquitin-dependent protein degradation and stress resistance in peripheral tissues including the intestine. A key regulator of this food response is the toll receptor domain protein TIR-1, which is dynamically regulated by direct binding between *tir-1* mRNA and *mir-71* (*mir-71/tir-1*) in AWC neurons. Absence of olfactory information (food odor) or abrogation of posttranscriptional regulation via *mir-71* results in elevated *tir-1* level. Chronic upregulation of *tir-1* in AWC neurons in *mir-71(n4115)* alters diet-evoked communication and blocks adjustments of intestinal proteostasis which correlates with decreased stress tolerance and lifespan.

Previous reports indicated that neuroendocrine signaling communicates proteotoxic stress between neurons and the intestine^24,25,35^. However, little information has been gathered on the physiological role of these systems. Given that proteostasis is linked to aging and longevity^1^, the olfactory system appears of particular importance since food-derived volatile cues in *D. melanogaster* and *C. elegans* affect lifespan^36–38^. Here, we provide new mechanistic insights showing that a neuronal olfactory circuit rewires proteolytic networks in intestinal cells, thus establishing an underlying concept for the regulation of food adaption (Fig. 5).

In this context, the primary AWC olfactory neurons are central in sensing and transducing food-derived information. Interestingly, AWCs are OFF-neurons which exhibit high activity (intracellular Ca^2+^ level) in the absence of food cues^29^, a condition that results in gradual upregulation of *tir-1* mRNA (Fig. 4i) and might suggest a functional relationship between neuronal activity and *tir-1* abundance. *tir-1* is also chronically upregulated in *mir-71(n4115)* (Fig. 4j), which indicates an aberrant neuroendocrine signaling that interferes with stress tolerance, aging, and organismal proteostasis. Since *tir-1* is a Ca^2+^ signaling scaffold protein located near synapse structures important for proper AWC development^20^, a similar function during adulthood in signal transduction can be hypothesized. However, in-depth activity studies in AWCs, connected primary sensory neurons, as well as interneurons will be needed to elucidate the exact role of TIR-1.

The TIR-1 ortholog sterile alpha and armadillo repeats containing protein 1 (SARM1) is predominantly expressed in the mammalian brain and facilitates clearance of damaged axons in response to trauma or disease, a process also known as Wallerian degeneration. Mechanistically, the SARM1-TIR domain provides NADase activity, which triggers depletion of the essential metabolite NAD^+^ and neuronal destruction^39^. Since changes in NAD^+^ levels are linked to proteasomal function, the NADase activity of SARM1 might directly affect ubiquitin-dependent protein degradation^40^. Here, the TIR domain from *C. elegans* TIR-1 provides a similar activity in axonal degeneration^41^. Interestingly, recent studies further associates SARM1 with the development of NAFLD (non-alcoholic fatty liver disease) induced by high fat diet^42^ as well as social interaction and cognitive flexibility in mice^43^. Together with our provided data on diet-dependent proteostasis and chemotactic behaviour this might indicate evolutionarily conserved concepts of TIR-1/SARM1.

The perception of food-related odors might coordinate food appreciation and selection with physiological and metabolic demands of the digestive tract. In this context, it is interesting to see that *E. coli* OP50 and HB101 differ in their nutrient composition; with higher carbohydrate level in HB101. Consequently, worms fed with HB101 show lower fat storage in comparison to OP50, indicative for diet-evoked metabolic changes^44^. Along this line, the sense of smell impacts the metabolic status especially in the context of obesity and fat storage in mice^45^ and regulates the production of digestive enzymes in humans^46^. Our observations show that the food source affects intestinal proteostasis^8^ (Fig. 3a, 4a, Supplementary Fig. 5a, b) which further suggests a direct link to the metabolic status of the organism. Together, the mechanistic insight on the organismal control via olfaction might open a new entry point to control and manipulate diet-related changes in proteostasis and metabolism with potential implication for associated diseases like obesity and diabetes.

## Materials and Methods

### *Caenorhabditis elegans* maintenance and transgenic lines

Nematodes were grown at 20°C (unless stated otherwise) on nematode growth medium (NGM) plates seeded with the bacterial *Escherichia coli* strain OP50 as a food source according to standard protocols and methods^47,48^. The N2 Bristol strain served as wild-type. All strains that were used in this study are listed in Supplementary Table 6. Transgenic strains expressing *Pnhx-2∷tir-1, Podr-3∷tir-1 and Pstr-1∷mir-71* were constructed via PCR amplification of N2 genomic DNA or cDNA (*nhx-2* promoter 602 bp genomic DNA^49^, *tir-1* 2793 bp cDNA, *tir-1 3’UTR* 587 bp cDNA, *odr-3* promoter 2755 bp genomic DNA^17^, *str-1* promoter 4000 bp genomic DNA^18^, premature *mir-71* 94 bp genomic DNA). Oligonucleotides used in this study are listed in Supplementary Table 7. PCR fragments were cloned into the IR101 (pDEST R4-R3) vector containing a *Prps0∷hygR∷mCherry* cassette for selection of transgenic animals. The constructs were particle bombarded into N2 hermaphrodites and selected as described previously^50^. Microparticle bombardment was done with the Bio-Rad Biolistic PDS-1000/HE with ¼” gap distance, 9 mm macrocarrier to screen distance, 28 inches of Hg vacuum and a 1350 p.s.i. rupture disc. Per bombardment, about 1 mg of 1 µm microcarrier gold beads were coated with 7 µg linearized DNA. Animals were allowed to recover for 1 h at room temperature and were then transferred to 90 mm enriched peptone plates seeded with *E. coli* C600 bacteria. After 3-4 days 100 µl hygromycin B (100 mg/ml), 600 µl Antibiotic-Antimycotic 1006 (GIBCO Life technologies) and kanamycin (final concentration 50 mg/ml) were added to each plate to select transgenic animals by hygromycin B resistance. About 10 days after bombardment viable animals were screened for mCherry fluorescence. Individuals positive for mCherry were singled and screened for homozygosity. Strains expressing the UFD substrate (UbV-GFP) were described previously^8^.

### CRISPR/Cas9 genome editing

CRISPR/Cas9 mutant alleles were generated as described^51^. Briefly, around 35-40 worms were injected with a mix containing 50 ng/μl P*eft-3:Cas9sv40:nls:tbb-2*^52^, 50 ng/μl plk111 (*U6prom:sgSRNA*)^53^, 50 ng/μl repair templates (see Supplementary Table 8) for homology directed repair, 3 ng/μl P*myo-2:mCherry* and 100 ng/μl of a plasmid expressing a neomycin resistance cassette (pBCN44 from^54^). Injected F0 were singled out and stable lines were selected on OP50-seeded NGM plates, containing (400 μg/ml) of neomycin (G-418). All surviving F1 hermaphrodites within each plate were pooled and tested by PCR for the desired event. In case of the *tir-1 3’UTR\** mutant the introduced mutations were spotted by restriction digestion of the PCR product with KpnI. Non-transgenic animals were cloned out to obtain homozygous isolates and the absence of the extrachromosomal array was verified by PCR. All alleles were validated by Sanger sequencing of the respective locus.

### Lifespan and survival assays

For lifespan assays 100 age-synchronized L4 larvae per strain were transferred to fresh NGM agar plates. Day 0 of the lifespan experiment refers to the first day after reaching adulthood. Within the first seven days animals were placed on new plates every day to separate them from F1s and to prevent starvation. After entering the post-reproductive phase, worms were only transferred when necessary. Survival was examined daily by checking pharyngeal pumping or avoidance behaviour in response to mechanical stimuli. All lifespan experiments were carried out in two biological replicates at 20°C. If indicated, NGM agar plates were supplemented with 25 μM fluorodeoxyuridine (FUdR) to prevent embryonic development and egg hatching^55^. Statistical details for all life span experiments are presented in Supplementary Table 1. To dissect sensitivity to endoplasmic reticulum (ER) stress conditions NGM plates were supplemented with 50 µg/µl tunicamycin (an equal amount of DMSO served as solvent control) followed by monitoring survival as described above. For proteasome inhibition NGM agar plates were supplemented with 10 μM bortezomib (BTZ) and L4 larvae were placed on the respective plates. Survival was determined for three consecutive days. Experiments were performed with 50 worms for three biological replicates. Heat stress was induced by incubation of age-synchronized L4 larvae at 32.5°C for 15 hr. Survival was analyzed in three biological replicates using 50-100 worms each.

### Western blotting

To prepare whole worm lysates, 100 animals were collected in 80 µl 1x SDS loading buffer. Subsequently, samples were boiled at 95°C for 5 min, sonicated (30 sec at 60% amplitude) and again boiled at 95°C for 5 min. Following, samples were centrifuged at 15,000 rpm for 5 min. For western blotting, protein lysates were resolved by SDS-PAGE using NuPAGE^®^ 4-12% Bis-Tris SDS-gels with the respective NuPAGE^®^ MES SDS running buffer (ThermoFischer Scientific; settings according to manufacturer’s instructions). A semi-dry blotting system (Bio-Rad, Trans-Blot Turbo) was applied with NuPAGE^®^ transfer buffer. Antibodies were diluted in 1x Roti^®^-Block (Carl Roth). The following antibodies were used in this study: Mouse Monoclonal anti-alpha Tubulin (Clone B-5-1-2) (Sigma-Aldrich; Cat# T6074 RRID:AB_477582), Living Colors^®^ A.v. Monoclonal Antibody (JL-8) (Mouse anti-GFP) (Clontech; Cat# 632380), Donkey anti-mouse IRDye® 800CW/680 (LI-COR; Cat# 926-32212 RRID:AB_621847). Visualization of fluorescent signals was achieved using the Odyssey scanner (LI-COR) and the Image Studio Lite v4.0 software.

### Microscopy

For fluorescent and DIC images the M165 FC stereomicroscope with a DFC 340 FX camera (Leica Camera) or the Axio Zoom V16 microscope with an Axiocam 506 mono camera (Carl Zeiss Microscopy GmbH) was used and images were processed with the Zen 2.3 pro software (Carl Zeiss Microscopy GmbH). For close-up images of the *C. elegans* head region, the Confocal Laser Scanning Microscope Zeiss Meta 710 (Carl Zeiss Microscopy GmbH) was used. To paralyze worms for imaging, animals were treated with 0.1 µg/ml levamisole (Sigma Aldrich) or 60 mM sodium azide (Carl Roth). For analyzing TIR-1∷GFP levels in AWC neurons age-synchronized L4 larvae of *tir-1∷gfp* worms, also expressing RFP under an AWC-specific promotor (*Podr-1∷rfp*), were grown until reaching day 1 of adulthood on OP50-seeded NGM plates supplemented with 25 µM BTZ or on plates supplemented with an equal volume of DMSO as solvent control, respectively. Worms were prepared as described above for confocal microscopy and obtained images were processed using ImageJ 1.52b. GFP fluorescence intensity was analyzed with the Imaris 9.l.2 software. Therefore, the volume of AWC neurons was determined in µm^3^ by creating a region of interest (ROI) based on RFP fluorescence with a threshold of 0.9 for background substraction. Within this ROI the GFP intensity was calculated as fluorescence intensity/µm^3^.

### RNA interference

RNAi was performed via the feeding method established for *C. elegans*^56,57^. Age-synchronized worms were transferred to RNAi plates seeded with *E. coli* HT115 bacteria expressing the respective double-stranded RNA (dsRNA). Clones carrying the vector against a gene of interest were either taken from the RNAi Collection (Ahringer) or the ORF-RNAi Resource (Vidal)^58^ (Source BioScience). As control condition, bacteria transformed with the empty pPD129.36 vector were used for feeding. To achieve an enhanced RNAi efficiency in neuronal tissues, additional knockdown of the RNaseT enzyme ERI-1 was performed as described^59^.

### RNA purification and qRT-PCR

Isolation of total RNA was performed using TRIzol (Invitrogen) and the QIAGEN RNeasy^®^ kit. Age-synchronized worms were washed two times with M9 buffer, resuspended in 1 ml TRIzol reagent and frozen at −80°C for at least 1 hr. Worms were homogenized by adding silica beads, followed by tissue disruption with the Precellys 24-Dual cell homogenizer (Peq-Lab). Next, 1-bromo-3-chloropropane was added to the samples followed by vigorous mixing and phase separation via centrifugation. The aqueous phase was used to isolate total RNA with the RNeasy^®^ Mini Kit (Qiagen) according to manufacturer’s instructions. cDNA synthesis was performed with 200 ng total RNA using the High-Capacity cDNA Reverse Transcription Kit (Applied Biosystems). Gene expression levels were measured via quantitative real time PCR (qRT-PCR) with the Brilliant III Ultra-Fast SYBR^®^ Green QPCR Master Mix (Agilent Technologies) and the Bio-Rad CFX96 Real-Time PCR Detection System. For each sample three technical replicates were analyzed and *cdc-42* and *pmp-3* were used for normalization^60^.

### Microarray analysis and RNA sequencing

Expression profiling of *mir-71* mutants versus WT was done using the Affymetrix^™^ *C. elegans* Gene Array 1.0. To this end, total RNA of around 200-300 age-synchronized day 1 adult hermaphrodites was extracted as described above. Microarray analysis was performed using four replicates per strain and was processed by the Cologne Center for Genomics (CCG, Cologne, Germany). Further quality control (analysis of RNA integrity number (RIN)) and gene expression profiling was performed. The obtained data were analyzed in detail with Partek^®^ Genomic Suite^®^ 6.6. Genome-wide transcriptional responses to proteotoxic stress were analyzed by RNA sequencing. Mid L4-staged N2 worms were put on OP50-seeded NGM plates containing 25 µM cadmium, 200 µM fluphenazine, 200 µM cantharidin or 10 µM and 25 µM BTZ, and then collected for total RNA extraction using the mirVana^™^ microRNA isolation kit (Ambion). Worms treated with *rpn-6.1* RNAi were put on RNAi plates from early L3 stage on and harvested as day 1 adults. For each condition triplicates of ∼2000 worms each were used for total RNA extraction. RNA quality control, library preparation of polyA-selected transcripts and subsequent Illumina^®^ sequencing was conducted by the CCG. Raw data were analyzed by the in-house Bioinformatics facility to compute fold changes relative to untreated N2 worms. Filtering of the data and heatmap generation was done using Microsoft Excel. During establishment of the conditions, the various treatments were analyzed for their capacity to block the proteasome and to induce autophagy. Proteotoxic stress was applied by either, fluphenazine and cantharidin treatment-induced autophagy, or by directly blocking the proteasome with 10 or 25 µM BTZ or *rpn-6.1* RNAi.

### Bacterial feeding and chemotaxis assays

To study the impact of different food sources on proteostasis, worms were grown on different bacterial strains and UFD substrate (UbV-GFP) or *tir-1* mRNA levels were monitored via western blotting and qRT-PCR, respectively. Here, age-synchronized L1 larvae were transferred to plates seeded with the respective *E. coli* strain. For inedible food experiments *E. coli* were treated with the antibiotic aztreonam (Az) that causes cell division stop and morphological changes inhibiting consequent ingestion by *C. elegans*^34,61^. OP50 were grown to log phase at 37°C, followed by Az treatment (final concentration 10 µg/ml) for additional 3 hr at 37°C and seeded to standard NGM plates 1 day before the experiment. OP50 expressing GFP (OP50-GFP) were used to control blocked ingestion after Az treatment (Fig. 4h). Age-synchronized worms were grown until day 1 of adulthood on *E.coli* OP50, transferred to Az containing plates, control plates or plates without food and cultivated for 6 hr. Following, animals were harvested for qRT-PCR as described above.

For studying chemotaxis of *C. elegans*, behavioural assays were adapted from S. Ward^33^. Chemotaxis studies were performed on freshly prepared Agar plates (2 % Agar; 5 mM KPO4, pH 6.0; 1 mM CaCl2; 1 mM MgSO4). Plates were separated in four equal quadrants (two test and two control quadrants). In each of the test quadrants 20 µl of OP50 bacteria containing 1 µM sodium azide were placed near the rim of the plate. LB medium was used as control condition.

Age-synchronized worms were grown on NGM plates seeded with OP50 bacteria until day 1 of adulthood, rinsed from plates with S-Basal medium and washed three times. Animals were then transferred to the centre of the chemotaxis plate. After the residual S-Basal medium was soaked in, the plates were sealed with parafilm and after a 1.5 hr incubation at room temperature the number of worms in each quadrant was counted. Individuals that did not cross the inner circle were excluded. The chemotaxis index (CI) was determined with the following equation:

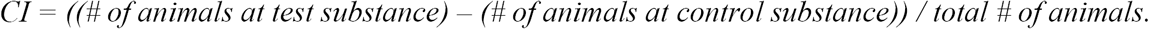

### Silencing of AWC olfactory neurons

For functional silencing of AWC activity^32^, synchronized worms carrying *Ex[ceh-36∷HisCl; myo-2∷GFP]* (gift from T. Ishihara)^62^ were grown until L4 larval stage either on *E. coli* OP50 or HB101, respectively. L4s are finally transferred to histamin (HA) containing NGM plates (10 mM final concentration) or control plates and allowed to grow until day 1 of adulthood, followed by assessment of proteostasis capability using the UFD substrate (UbV-GFP) and according western blotting.

### Statistical analysis

For statistical analysis the GraphPad Prism 5 software was used. A two-tailed paired Student’s t-test or one-way ANOVA with post-hoc test were used to analyze statistical significance of mRNA levels, fluorescence intensity in AWC neurons, chemotaxis, and proteotoxic stress assays. Results are given as mean and standard error of the mean (SEM). For survival assays, significance was determined using the Log-rank (Mantel-Cox) test.

## Acknowledgments

We thank Y. Kohara, A. Segref and the *Caenorhabditis* Genetics Center (funded by the NIH National Center for Research Resources), the Dana-Farber Cancer Institute, Addgene and Geneservice Ltd for plasmids, cDNAs, and strains. We thank the CECAD Imaging facility for support with confocal microscopy and the Cologne Center for Genomics for microarray analysis and RNA sequencing.

---

**Supplementary Figure 1.**
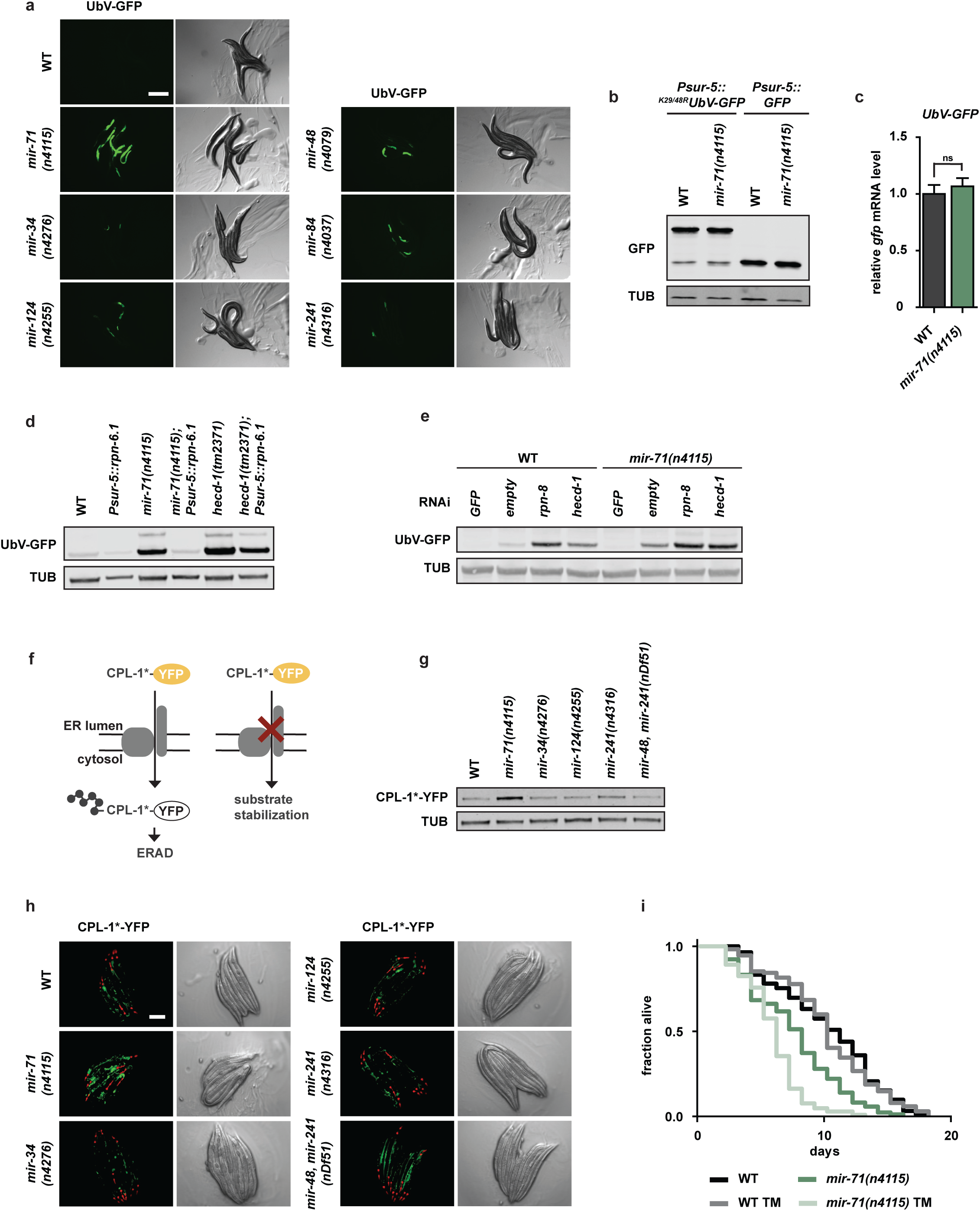
| Functional proteostasis requires the microRNA *mir-71*. **a**, The *mir-71(n4115)* deletion allele exhibits stabilization of the UFD substrate. Representative fluorescent images of day 1 adult worms with indicated genotypes. Scale bar: 300 µm. **b**, Accumulation of the UFD substrate *Psur-5∷UbV-GFP* in *mir-71(n4115)* is not a consequence of changes in gene expression. Representative western blot of worm lysates of indicated genotypes detect UbV-GFP, GFP, and tubulin (TUB). **c**, *UbV-GFP* mRNA levels are not increased in *mir-71(n4155)* mutant worms as tested by qRT-PCR (mean values ± SEM obtained from n=3 technical replicates; ns=not significant). **d**, *mir-71* affects UFD substrate degradation downstream of ubiquitylation. Western blot from worm lysates with indicated genotypes showing UbV-GFP and tubulin (TUB) levels. **e**, Inhibition of proteasomal degradation in *mir-71(n4115)* worms further elevates UFD substrate levels. Representative western blot depicting UbV-GFP and tubulin (TUB) in RNAi-depleted worms lacking *rpn-8* or *hecd-1*. **f**, The CPL-1*-YFP model substrate for studying ER-associated protein degradation (ERAD). **g**, The *mir-71(n4115)* deletion mutant accumulates the ERAD substrate. Western blot of worms imaged in (**h**) detecting CPL-1*-YFP and tubulin (TUB). **h**, Fluorescent images of day 1 adult worms expressing the ERAD substrate (CPL-1*-YFP); pharyngeal expression of *Pmyo-2∷mCherry* served as transgenic marker. Scale bar: 250 µm. **i**, *mir-71(n4115)* exhibits increased sensitivity to ER stress induced by tunicamycin (TM) treatment (50 µg/µl). For statistics details see Supplementary Table 1.

**Supplementary Figure 2.**
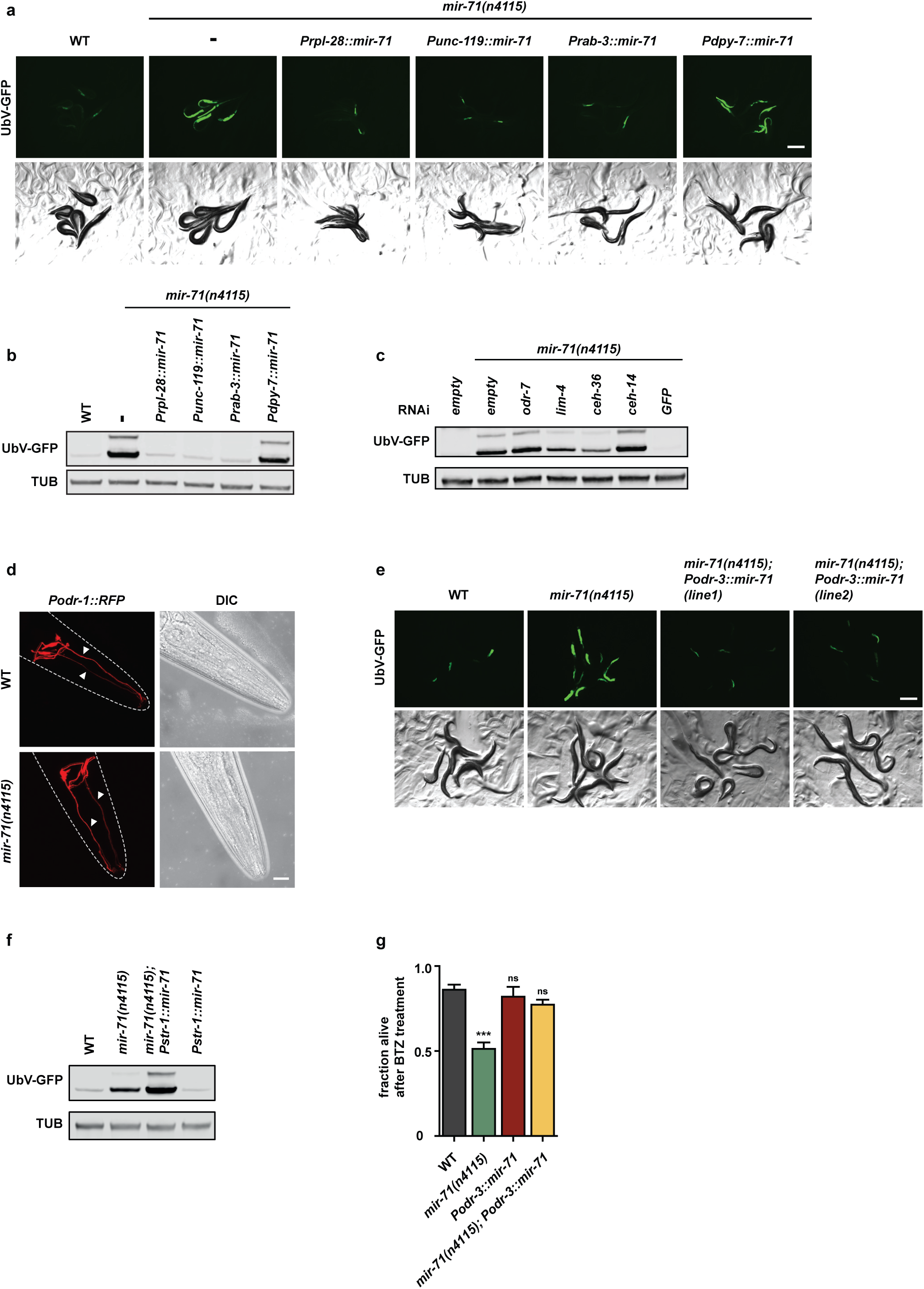
| Neuron-specific expression of *mir-71* modulates proteostasis. **a**, Neuronal expression of *mir-71* rescues protein degradation defects of *mir-71(n4115)*. Representative fluorescent images of day 1 adult *mir-71(n4115)* worms expressing the UFD substrate and the indicated tissue-specific rescue constructs *Prpl-28∷mir-71* (ubiquitous), *Punc-119∷mir-71* and *Prab-3∷mir-71* (both pan-neuronal), *Pdpy7∷mir-71* (hypodermal). Scale bar: 300 µm. **b**, Representative western blot of worms imaged in (**a**) detect UbV-GFP and tubulin (TUB). **c**, Inhibition of neuronal development ameliorates *mir-71(n4115)* proteolytic defects. Western blot of *mir-71(n4115)* worms treated with RNAi against the indicated transcription factors show UbV-GFP and tubulin (TUB) levels. **d**, Deletion of *mir-71* does not affect AWC neuron morphology. Confocal microscopy images show AWC neurons (*Podr-1∷RFP*) and differential interference contrast images (DIC) of the *C. elegans* head region. Scale bar: 15 µm. **e**, AWC-specific rescue of the *mir-71(n4115)* deletion mutant (*mir-71(n4115); Podr-3∷mir-71*) restores protein degradation. Representative fluorescent images of day 1 adult worms expressing the UFD substrate. Scale bar: 300 µm. **f**, AWB-specific expression of *mir-71* (*Pstr-1∷mir-71*) does not rescue proteostasis defects of *mir-71(n4115)* animals. Representative western blot of worm lysates with the indicated genotypes show UbV-GFP and tubulin (TUB). **g**, AWC-specific expression of *mir-71* increases survival upon proteasome inhibition (BTZ). Data represent mean values ± SEM generated from n=3 independent experiments using at least 50 worms; ***p < 0.001, ns=not significant.

**Supplementary Figure 3.**
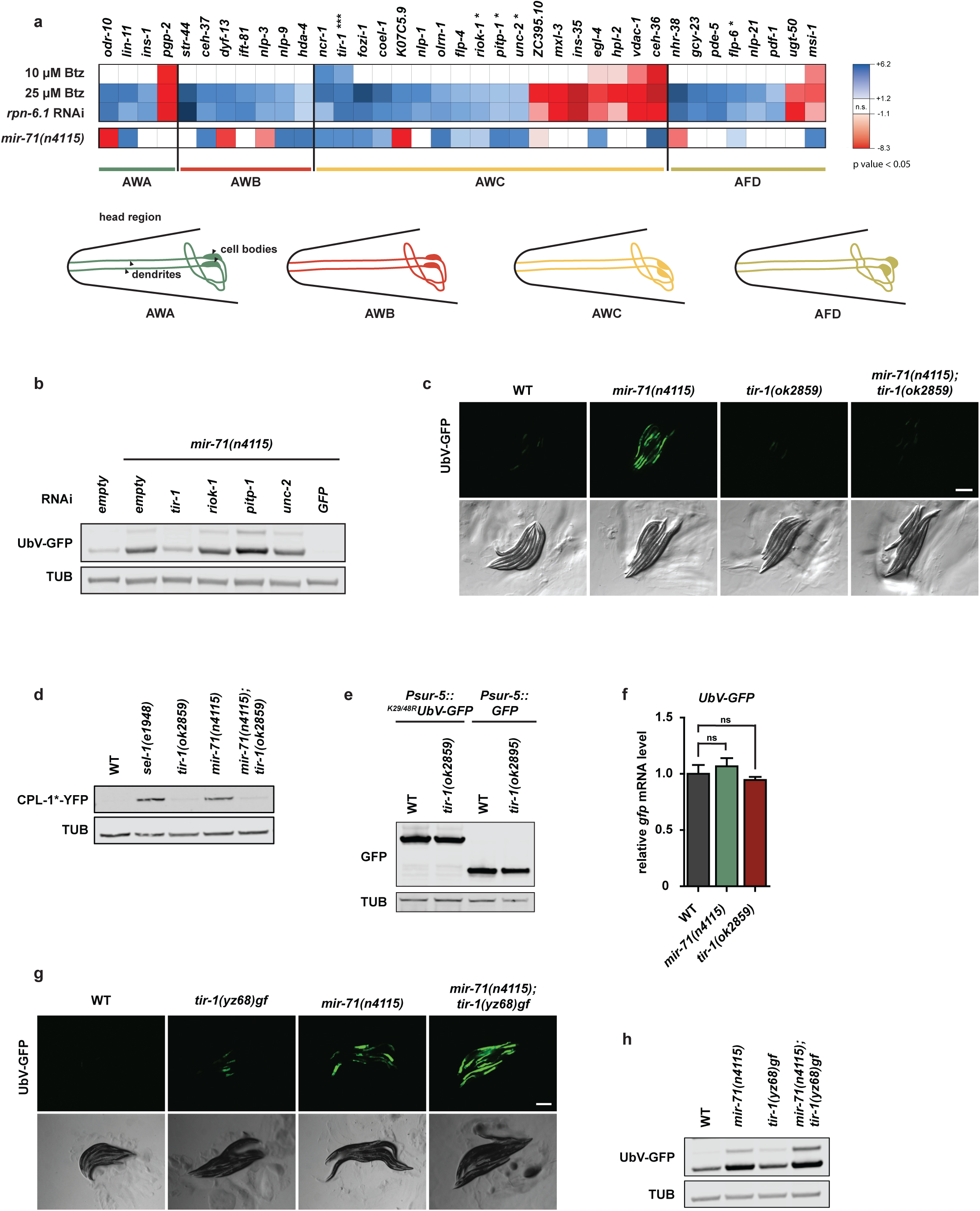
| Genetic interaction between *mir-71* and *tir-1* affects proteostasis. **a**, *tir-1* mRNA levels are upregulated upon induction of proteotoxic stress and in *mir-71(n4115)*. Heat map of mRNA sequencing data for factors present in different chemosensory neurons. A schematic depiction of the investigated amphid sensillum neurons in the *C. elegans* head region is shown at the bottom. Olfactory neurons: AWA = Amphid wing A cell (green), AWB = Amphid wing B cells (red), AWC = Amphid wing C cells (yellow); thermosensory neuron: AFD = Amphid neurons with finger-like (AFD) ciliated endings (light green). mRNA level in wild-type (WT) worms with proteasomal inhibition (10 µM or 25 µM BTZ or *rpn-6.1* RNAi) and in *mir-71(n4115)* deletion mutants are compared to WT (untreated) worms. Asterisks (*) mark candidate transcripts that contain one (*) or up to three (***) potential *mir-71* binding sites; n.s.=not significant. **b**, *tir-1* depletion rescues *mir-71(n4115)* proteolytic defects. Western blot of *mir-71(n4115)* worm lysates treated with RNAi against the indicated factors show UbV-GFP and tubulin (TUB) levels. **c**, The *tir-1(ok2859)* deletion allele suppresses *mir-71(n4115)*-induced proteolytic defects. Representative fluorescent images of day 1 adult worms expressing the UFD substrate. Scale bar: 300 µm. **d**, *tir-1(ok2859)* deletion is able to suppress ERAD substrate (CPL-1*-YFP) stabilization in *mir-71(n4115)* worms. Representative western blot of worm lysates with indicated genotypes, detect CPL-1*-YFP and tubulin (TUB). **e**, UFD substrate stabilization in the *tir-1(ok2859)* deletion mutant is not caused by changes in gene expression. Western blots from worm lysates show UbV-GFP, GFP, and tubulin (TUB). **f**, *UbV-GFP* transcript levels are not altered by *mir-71* or *tir-1* deletions as detected by qRT-PCR. Data are shown as mean values ± SEM obtained from n=3 technical replicates; ns=not significant. **g-h**, The *tir-1* gain-of-function allele *tir-1(yz68)gf* leads to mild UPS defects. **g**, Representative fluorescent images of day 1 adult worms expressing the UFD substrate. Scale bar: 300 µm. **h**, Detection of the GFP signals shown in (**g**) via western blot, showing UbV-GFP and tubulin (TUB) level.

**Supplementary Figure 4.**
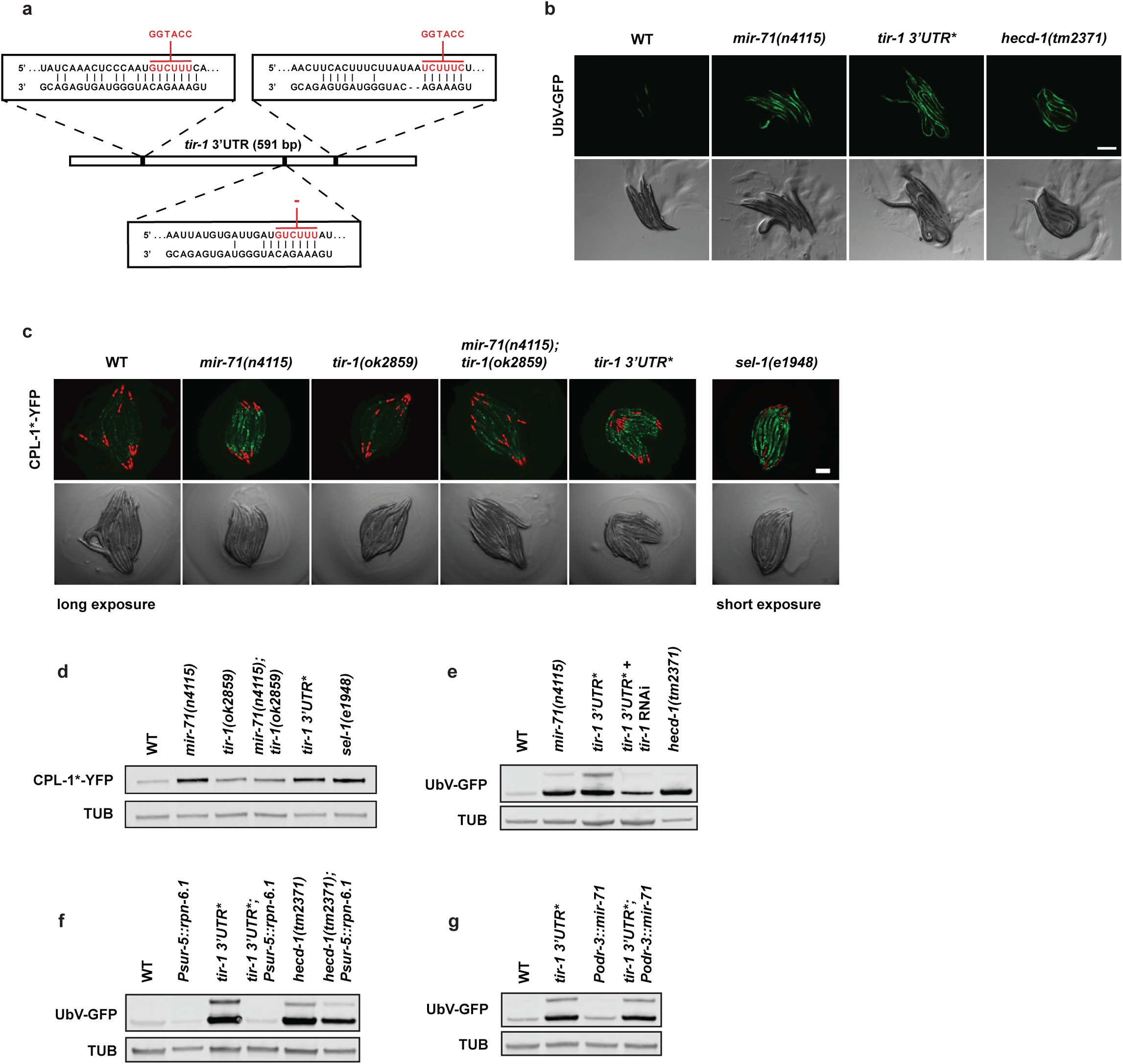
| Loss of *mir-71* binding sites in *tir-1* 3’UTR resembles proteolytic defects. **a**, Schematic overview of the *tir-1 3’UTR* sequence with indicated base pairs (bp). Mutated *mir-71* binding sites in *tir-1 3’UTR\** are shown in red. **b**, The *tir-1 3’UTR\** mutation induces proteolytic defects. Representative fluorescent images of day 1 adult worms expressing the UFD substrate. Scale bar: 300 µm. **c**, The *tir-1 3’UTR\** mutant accumulates the ERAD substrate. Fluorescent images of day 1 adult worms expressing the ERAD substrate (CPL-1*-YFP); pharyngeal expression of *Pmyo-2∷mCherry* served as transgenic marker. Scale bar: 250 µm. **d-g**, Representative western blots from worm lysates of indicated genotypes detect CPL-1*-YFP (**d**) UbV-GFP (**e-f**) and tubulin (TUB) protein level. **d**, The *tir-1 3’UTR\** mutation affects ERAD. Detection of the fluorescent signals shown in (**c**). **e**, RNAi-mediated *tir-1* depletion rescues UFD substrate stabilization in the *tir-1 3’UTR\** mutant. **f**, The *tir-1 3’UTR\** mutant triggers proteostasis defects downstream of substrate ubiquitylation. **g**, AWC-specific expression of *mir-71* does not suppress proteostasis defects caused by the *tir-1 3’UTR\** mutation.

**Supplementary Figure 5.**
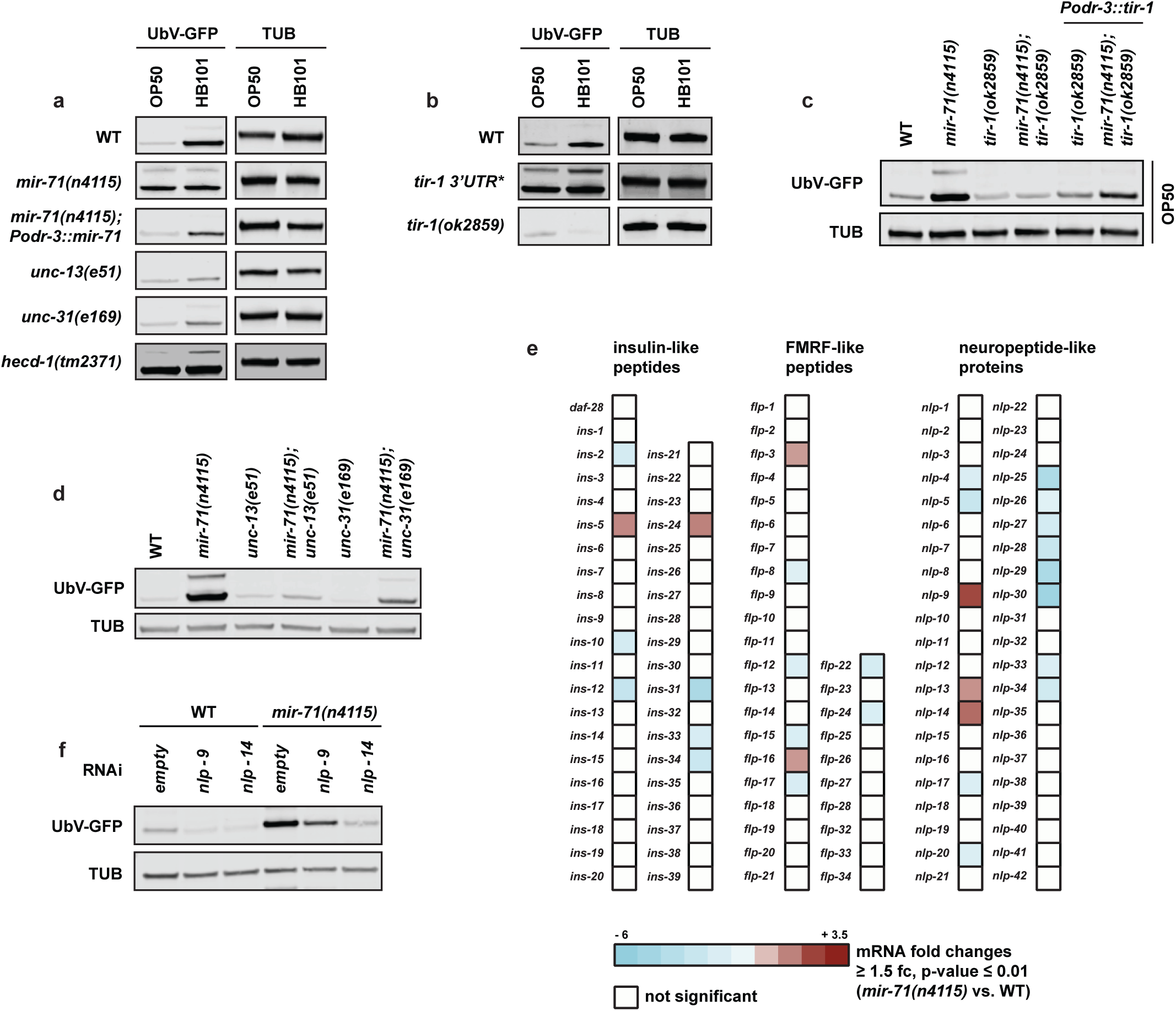
| Neuronal signaling facilitates *mir-71/tir-1* regulation of proteostasis. **a-b**, Different food sources influence organismal protein degradation through *mir-71/tir-1* regulation. Animals were grown on indicated bacteria prior to lysis for western blotting. Representative western blots from day 1 adult worms of indicated genotypes show UbV-GFP and tubulin (TUB) level. **c**, AWC-specific *tir-1* expression is important for food perception and proteostasis. Animals were grown on HB101 bacteria prior to worm lysis. Western blot of worm lysates with the indicated genotypes detecting UbV-GFP and tubulin (TUB) level. **d**, Inhibition of neuronal signal transduction suppresses *mir-71(n4115)* proteolytic defects. Western blot from worm lysates with indicated genotypes show UbV-GFP and tubulin (TUB) protein levels. **e**, Microarray data of *mir-71(n4115)* compared to wild-type (WT) reveals misregulation of neuropeptide expression. **f**, NLP-9 and NLP-14 neuropeptides are important for proteostasis signaling of the *mir-71(n4115)* mutant. Representative western blot from worms treated with RNAi against the indicated factors detect UbV-GFP and tubulin (TUB) levels.

**Supplementary Table 1.**
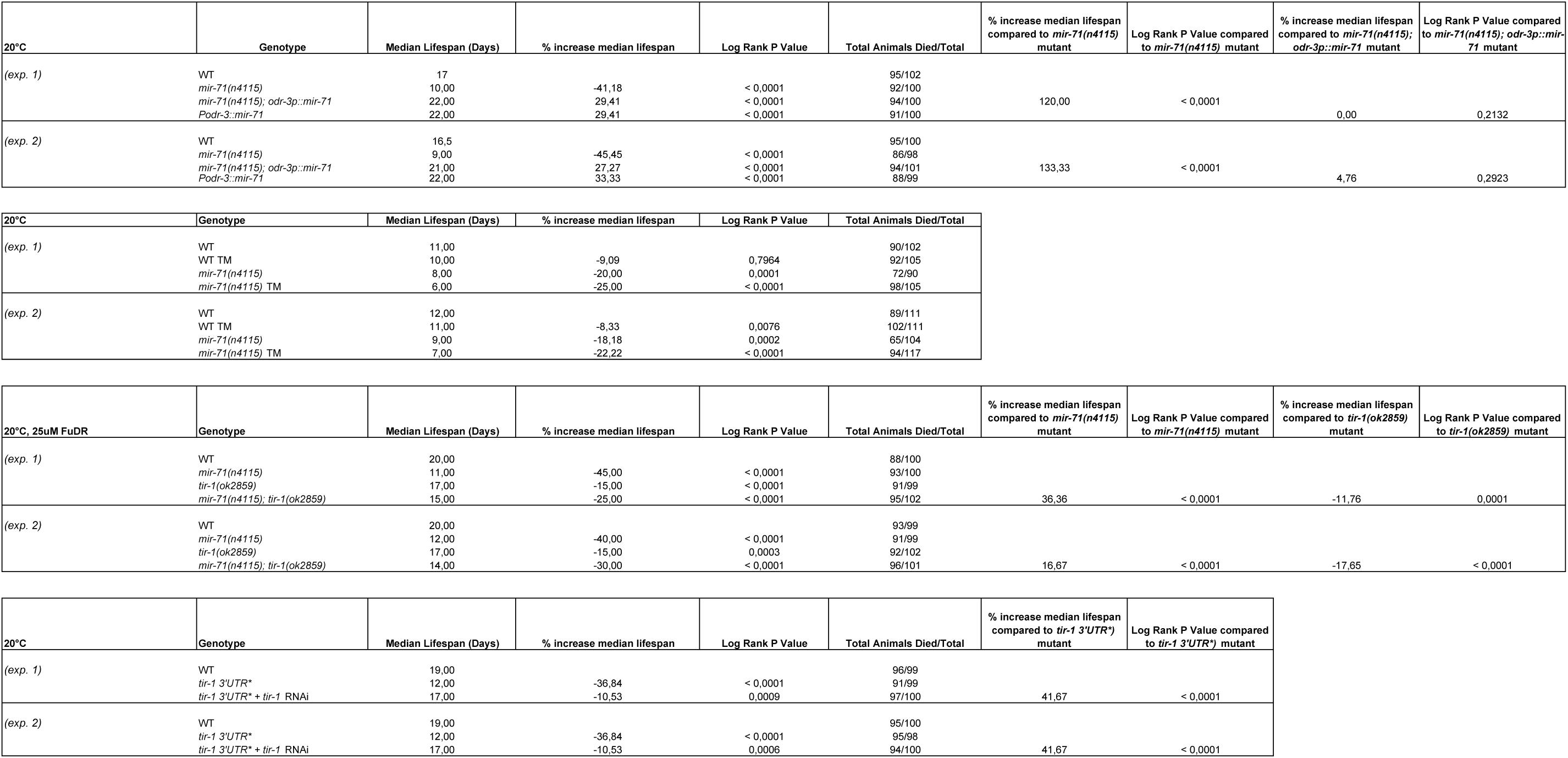
| Summary of lifespan experiments.

**Supplementary Table 2.**
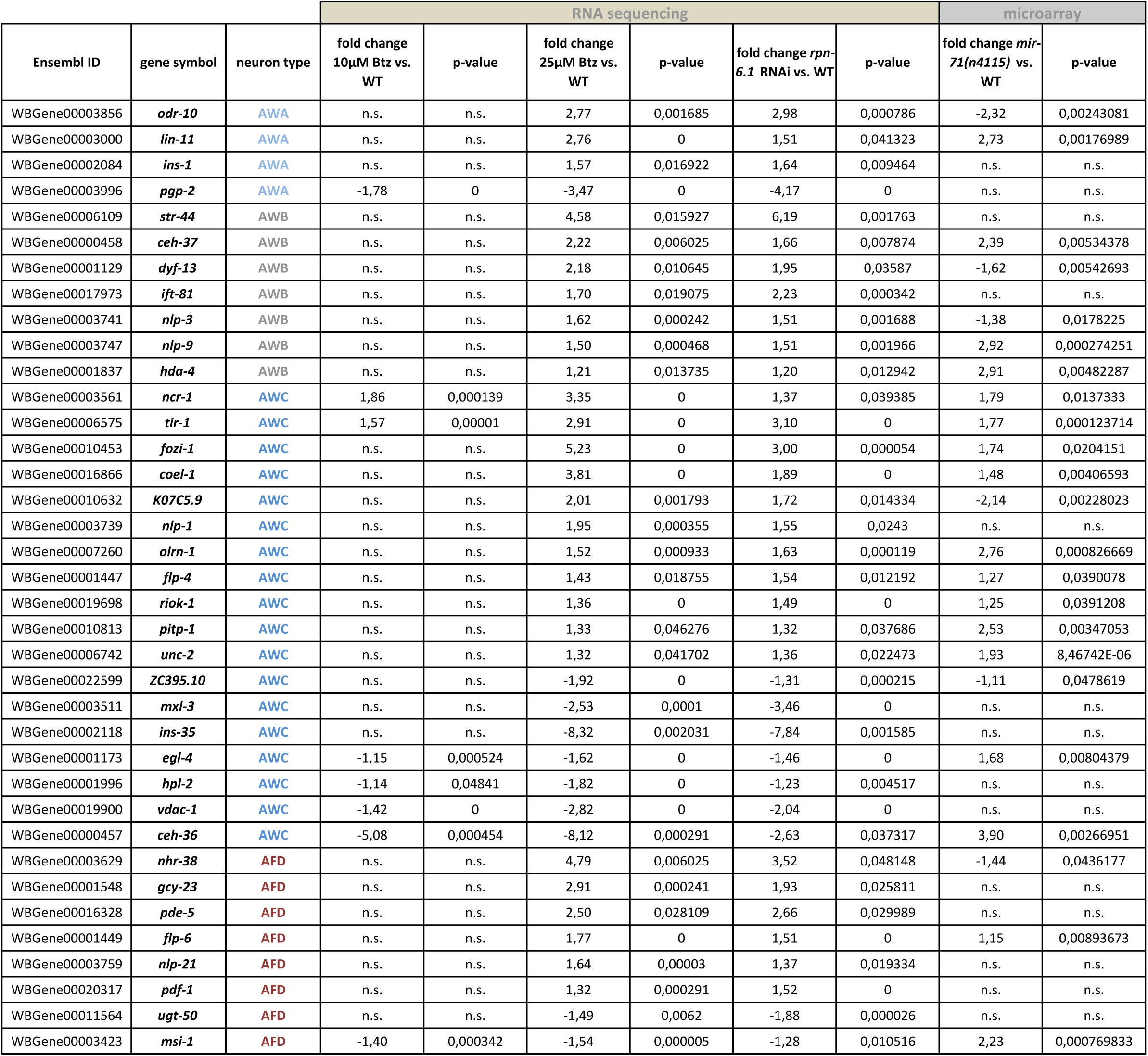
| mRNAs regulated in amphid sensory neurons upon proteasomal stress and *mir-71(n4115)* deletion.

**Supplementary Table 3.**
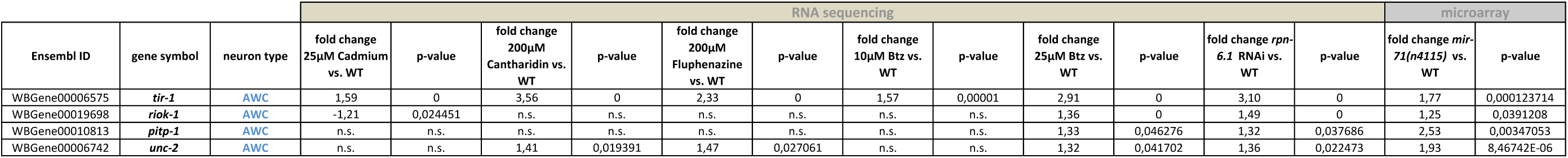
| mRNAs regulated in AWC neurons upon proteasomal stress and *mir-71(n4115)* deletion.

**Supplementary Table 4.** | Neuropeptide mRNAs regulated upon *mir-71(n4115)* deletion.

insulin-like peptides
| gene symbol | fold change(*mir-71(n4115)* vs. WT) | p-value |
| --- | --- | --- |
| *ins-2* | -2,2646 | 0,004975 |
| *ins-5* | 1,8976 | 0,001525 |
| *ins-10* | -2,2401 | 0,005381 |
| *ins-12* | -3,1031 | 0,000113 |
| *ins-24* | 2,0026 | 0,000179 |
| *ins-31* | -4,6013 | 0,003063 |
| *ins-33* | -2,0609 | 0,002678 |
| *ins-34* | -2,8311 | 0,000204 |

FRMF-like peptides
| gene symbol | fold change(*mir-71(n4115)* vs. WT) | p-value |
| --- | --- | --- |
| *flp-3* | 1,5704 | 0,000391 |
| *flp-8* | -1,6706 | 0,007151 |
| *flp-12* | -1,8135 | 0,003158 |
| *flp-15* | -2,0221 | 0,003966 |
| *flp-16* | 1,6139 | 0,002402 |
| *flp-17* | -2,0482 | 0,002479 |
| *flp-22* | -2,2135 | 0,003794 |
| *flp-24* | -2,0124 | 0,006178 |

neuropeptide-like proteins
| gene symbol | fold change(*mir-71(n4115)* vs. WT) | p-value |
| --- | --- | --- |
| *nlp-4* | -1,9161 | 0,000829 |
| *nlp-5* | -2,9974 | 0,000056 |
| *nlp-9* | 2,9161 | 0,000274 |
| *nlp-13* | 1,7180 | 0,001724 |
| *nlp-14* | 2,3555 | 0,000402 |
| *nlp-17* | -1,8117 | 0,001413 |
| *nlp-20* | -1,8869 | 0,009217 |
| *nlp-25* | -4,4854 | 0,000144 |
| *nlp-26* | -1,5791 | 0,006250 |
| *nlp-27* | -2,3196 | 0,001470 |
| *nlp-28* | -3,0186 | 0,000266 |
| *nlp-29* | -4,5787 | 0,002695 |
| *nlp-30* | -5,2101 | 0,006844 |
| *nlp-33* | -1,7984 | 0,000068 |
| *nlp-34* | -2,5120 | 0,006185 |
| *nlp-40* | -1,6439 | 0,000759 |

Affymetrix EleGene-1\_0-st-v1.na34.ce6
≥ 1.5 fold change, p-value ≤ 0,01

**Supplementary Table 5.**
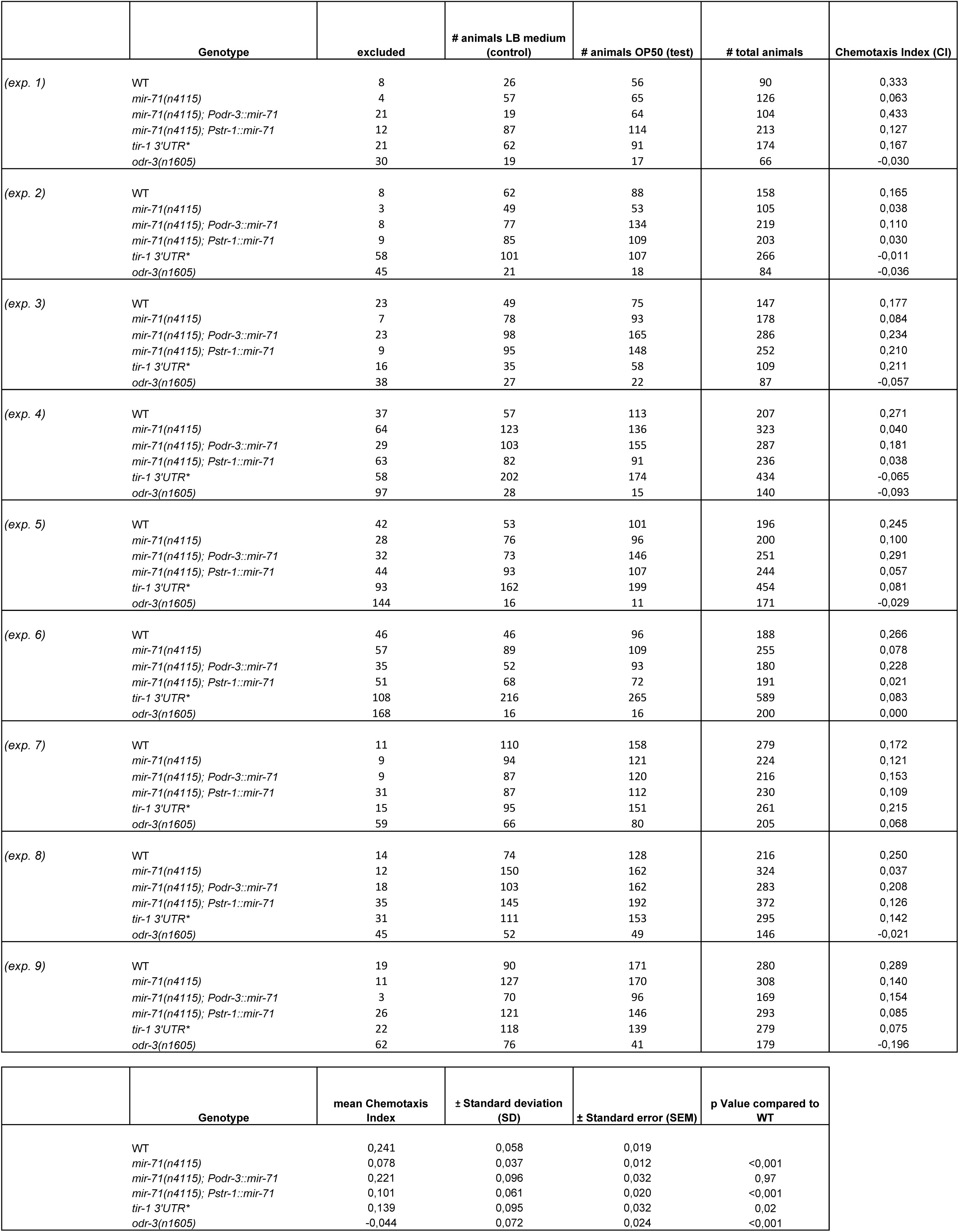
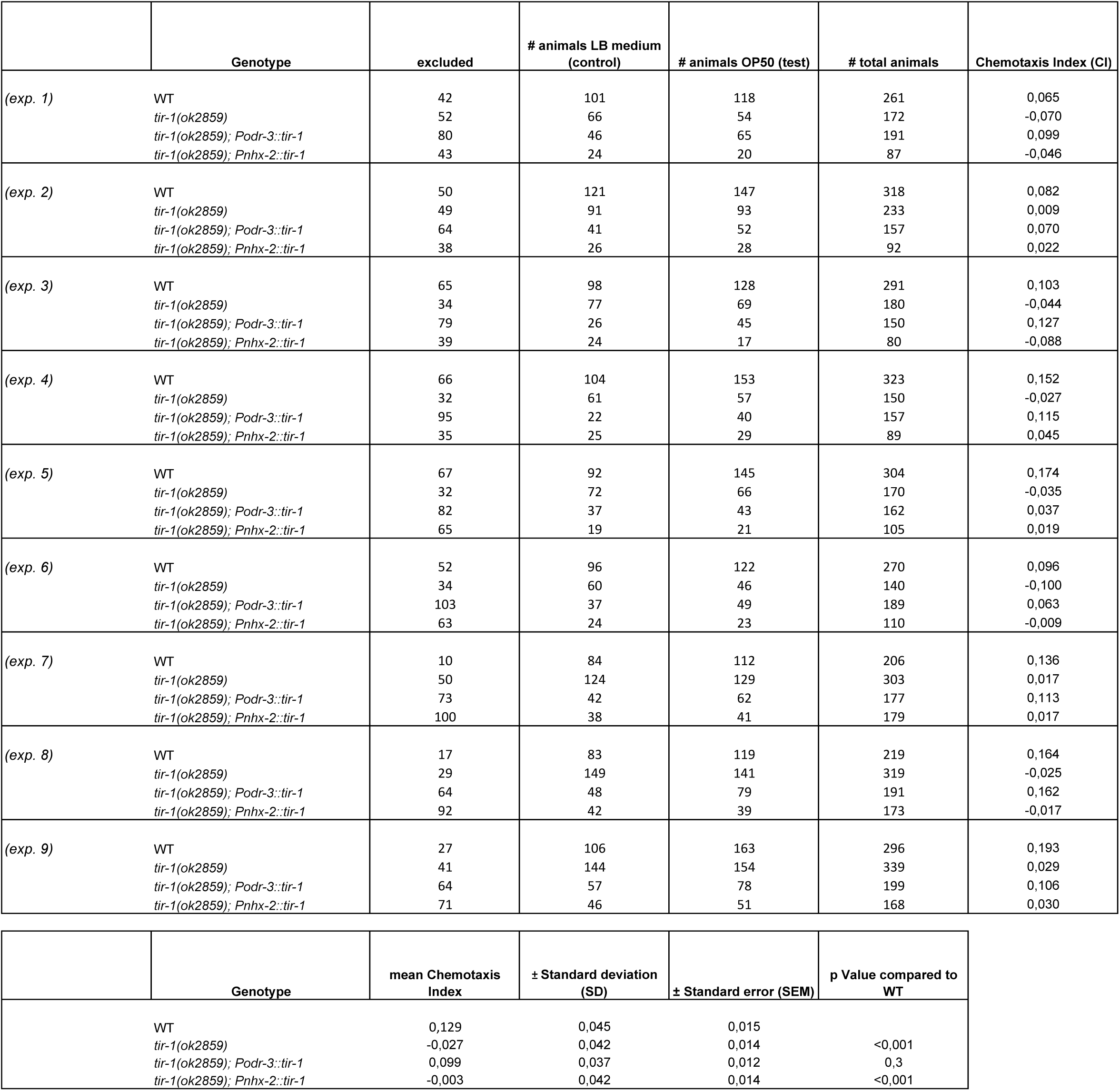
| Summary of chemotaxis experiments.

|  | Genotype | excluded | # animals LB medium (control) | # animals OP50 (test) | # total animals | Chemotaxis Index (CI) |
| --- | --- | --- | --- | --- | --- | --- |
| (exp. 1) | WT | 8 | 26 | 56 | 90 | 0,333 |
|  | *mir-71(n4115)* | 4 | 57 | 65 | 126 | 0,063 |
|  | *mir-71(n4115); Podr-3::mir-71* | 21 | 19 | 64 | 104 | 0,433 |
|  | *mir-71(n4115); Pstr-1::mir-71* | 12 | 87 | 114 | 213 | 0,127 |
|  | *tir-1 3'UTR\** | 21 | 62 | 91 | 174 | 0,167 |
|  | *odr-3(n1605)* | 30 | 19 | 17 | 66 | -0,030 |
| (exp. 2) | WT | 8 | 62 | 88 | 158 | 0,165 |
|  | *mir-71(n4115)* | 3 | 49 | 53 | 105 | 0,038 |
|  | *mir-71(n4115); Podr-3::mir-71* | 8 | 77 | 134 | 219 | 0,110 |
|  | *mir-71(n4115); Pstr-1::mir-71* | 9 | 85 | 109 | 203 | 0,030 |
|  | *tir-1 3'UTR\** | 58 | 101 | 107 | 266 | -0,011 |
|  | *odr-3(n1605)* | 45 | 21 | 18 | 84 | -0,036 |
| (exp. 3) | WT | 23 | 49 | 75 | 147 | 0,177 |
|  | *mir-71(n4115)* | 7 | 78 | 93 | 178 | 0,084 |
|  | *mir-71(n4115); Podr-3::mir-71* | 23 | 98 | 165 | 286 | 0,234 |
|  | *mir-71(n4115); Pstr-1::mir-71* | 9 | 95 | 148 | 252 | 0,210 |
|  | *tir-1 3'UTR\** | 16 | 35 | 58 | 109 | 0,211 |
|  | *odr-3(n1605)* | 38 | 27 | 22 | 87 | -0,057 |
| (exp. 4) | WT | 37 | 57 | 113 | 207 | 0,271 |
|  | *mir-71(n4115)* | 64 | 123 | 136 | 323 | 0,040 |
|  | *mir-71(n4115); Podr-3::mir-71* | 29 | 103 | 155 | 287 | 0,181 |
|  | *mir-71(n4115); Pstr-1::mir-71* | 63 | 82 | 91 | 236 | 0,038 |
|  | *tir-1 3'UTR\** | 58 | 202 | 174 | 434 | -0,065 |
|  | *odr-3(n1605)* | 97 | 28 | 15 | 140 | -0,093 |
| (exp. 5) | WT | 42 | 53 | 101 | 196 | 0,245 |
|  | *mir-71(n4115)* | 28 | 76 | 96 | 200 | 0,100 |
|  | *mir-71(n4115); Podr-3::mir-71* | 32 | 73 | 146 | 251 | 0,291 |
|  | *mir-71(n4115); Pstr-1::mir-71* | 44 | 93 | 107 | 244 | 0,057 |
|  | *tir-1 3'UTR\** | 93 | 162 | 199 | 454 | 0,081 |
|  | *odr-3(n1605)* | 144 | 16 | 11 | 171 | -0,029 |
| (exp. 6) | WT | 46 | 46 | 96 | 188 | 0,266 |
|  | *mir-71(n4115)* | 57 | 89 | 109 | 255 | 0,078 |
|  | *mir-71(n4115); Podr-3::mir-71* | 35 | 52 | 93 | 180 | 0,228 |
|  | *mir-71(n4115); Pstr-1::mir-71* | 51 | 68 | 72 | 191 | 0,021 |
|  | *tir-1 3'UTR\** | 108 | 216 | 265 | 589 | 0,083 |
|  | *odr-3(n1605)* | 168 | 16 | 16 | 200 | 0,000 |
| (exp. 7) | WT | 11 | 110 | 158 | 279 | 0,172 |
|  | *mir-71(n4115)* | 9 | 94 | 121 | 224 | 0,121 |
|  | *mir-71(n4115); Podr-3::mir-71* | 9 | 87 | 120 | 216 | 0,153 |
|  | *mir-71(n4115); Pstr-1::mir-71* | 31 | 87 | 112 | 230 | 0,109 |
|  | *tir-1 3'UTR\** | 15 | 95 | 151 | 261 | 0,215 |
|  | *odr-3(n1605)* | 59 | 66 | 80 | 205 | 0,068 |
| (exp. 8) | WT | 14 | 74 | 128 | 216 | 0,250 |
|  | *mir-71(n4115)* | 12 | 150 | 162 | 324 | 0,037 |
|  | *mir-71(n4115); Podr-3::mir-71* | 18 | 103 | 162 | 283 | 0,208 |
|  | *mir-71(n4115); Pstr-1::mir-71* | 35 | 145 | 192 | 372 | 0,126 |
|  | *tir-1 3'UTR\** | 31 | 111 | 153 | 295 | 0,142 |
|  | *odr-3(n1605)* | 45 | 52 | 49 | 146 | -0,021 |
| (exp. 9) | WT | 19 | 90 | 171 | 280 | 0,289 |
|  | *mir-71(n4115)* | 11 | 127 | 170 | 308 | 0,140 |
|  | *mir-71(n4115); Podr-3::mir-71* | 3 | 70 | 96 | 169 | 0,154 |
|  | *mir-71(n4115); Pstr-1::mir-71* | 26 | 121 | 146 | 293 | 0,085 |
|  | *tir-1 3'UTR\** | 22 | 118 | 139 | 279 | 0,075 |
|  | *odr-3(n1605)* | 62 | 76 | 41 | 179 | -0,196 |

|  | Genotype | mean Chemotaxis Index | ± Standard deviation (SD) | ± Standard error (SEM) | p Value compared to WT |
| --- | --- | --- | --- | --- | --- |
|  | WT | 0,241 | 0,058 | 0,019 |  |
|  | *mir-71(n4115)* | 0,078 | 0,037 | 0,012 | <0,001 |
|  | *mir-71(n4115); Podr-3::mir-71* | 0,221 | 0,096 | 0,032 | 0,97 |
|  | *mir-71(n4115); Pstr-1::mir-71* | 0,101 | 0,061 | 0,020 | <0,001 |
|  | *tir-1 3'UTR\** | 0,139 | 0,095 | 0,032 | 0,02 |
|  | *odr-3(n1605)* | -0,044 | 0,072 | 0,024 | <0,001 |
| (exp. 1) | WT | 42 | 101 | 118 | 261 | 0,065 |
|  | *tir-1(ok2859)* | 52 | 66 | 54 | 172 | -0,070 |
|  | *tir-1(ok2859); Podr-3::tir-1* | 80 | 46 | 65 | 191 | 0,099 |
|  | *tir-1(ok2859); Pnhx-2::tir-1* | 43 | 24 | 20 | 87 | -0,046 |
| (exp. 2) | WT | 50 | 121 | 147 | 318 | 0,082 |
|  | *tir-1(ok2859)* | 49 | 91 | 93 | 233 | 0,009 |
|  | *tir-1(ok2859); Podr-3::tir-1* | 64 | 41 | 52 | 157 | 0,070 |
|  | *tir-1(ok2859); Pnhx-2::tir-1* | 38 | 26 | 28 | 92 | 0,022 |
| (exp. 3) | WT | 65 | 98 | 128 | 291 | 0,103 |
|  | *tir-1(ok2859)* | 34 | 77 | 69 | 180 | -0,044 |
|  | *tir-1(ok2859); Podr-3::tir-1* | 79 | 26 | 45 | 150 | 0,127 |
|  | *tir-1(ok2859); Pnhx-2::tir-1* | 39 | 24 | 17 | 80 | -0,088 |
| (exp. 4) | WT | 66 | 104 | 153 | 323 | 0,152 |
|  | *tir-1(ok2859)* | 32 | 61 | 57 | 150 | -0,027 |
|  | *tir-1(ok2859); Podr-3::tir-1* | 95 | 22 | 40 | 157 | 0,115 |
|  | *tir-1(ok2859); Pnhx-2::tir-1* | 35 | 25 | 29 | 89 | 0,045 |
| (exp. 5) | WT | 67 | 92 | 145 | 304 | 0,174 |
|  | *tir-1(ok2859)* | 32 | 72 | 66 | 170 | -0,035 |
|  | *tir-1(ok2859); Podr-3::tir-1* | 82 | 37 | 43 | 162 | 0,037 |
|  | *tir-1(ok2859); Pnhx-2::tir-1* | 65 | 19 | 21 | 105 | 0,019 |
| (exp. 6) | WT | 52 | 96 | 122 | 270 | 0,096 |
|  | *tir-1(ok2859)* | 34 | 60 | 46 | 140 | -0,100 |
|  | *tir-1(ok2859); Podr-3::tir-1* | 103 | 37 | 49 | 189 | 0,063 |
|  | *tir-1(ok2859); Pnhx-2::tir-1* | 63 | 24 | 23 | 110 | -0,009 |
| (exp. 7) | WT | 10 | 84 | 112 | 206 | 0,136 |
|  | *tir-1(ok2859)* | 50 | 124 | 129 | 303 | 0,017 |
|  | *tir-1(ok2859); Podr-3::tir-1* | 73 | 42 | 62 | 177 | 0,113 |
|  | *tir-1(ok2859); Pnhx-2::tir-1* | 100 | 38 | 41 | 179 | 0,017 |
| (exp. 8) | WT | 17 | 83 | 119 | 219 | 0,164 |
|  | *tir-1(ok2859)* | 29 | 149 | 141 | 319 | -0,025 |
|  | *tir-1(ok2859); Podr-3::tir-1* | 64 | 48 | 79 | 191 | 0,162 |
|  | *tir-1(ok2859); Pnhx-2::tir-1* | 92 | 42 | 39 | 173 | -0,017 |
| (exp. 9) | WT | 27 | 106 | 163 | 296 | 0,193 |
|  | *tir-1(ok2859)* | 41 | 144 | 154 | 339 | 0,029 |
|  | *tir-1(ok2859); Podr-3::tir-1* | 64 | 57 | 78 | 199 | 0,106 |
|  | *tir-1(ok2859); Pnhx-2::tir-1* | 71 | 46 | 51 | 168 | 0,030 |

|  | Genotype | mean Chemotaxis Index | ± Standard deviation (SD) | ± Standard error (SEM) | p Value compared to WT |
| --- | --- | --- | --- | --- | --- |
|  | WT | 0,129 | 0,045 | 0,015 |  |
|  | *tir-1(ok2859)* | -0,027 | 0,042 | 0,014 | <0,001 |
|  | *tir-1(ok2859); Podr-3::tir-1* | 0,099 | 0,037 | 0,012 | 0,3 |
|  | *tir-1(ok2859); Pnhx-2::tir-1* | -0,003 | 0,042 | 0,014 | <0,001 |

**Supplementary Table 6.**
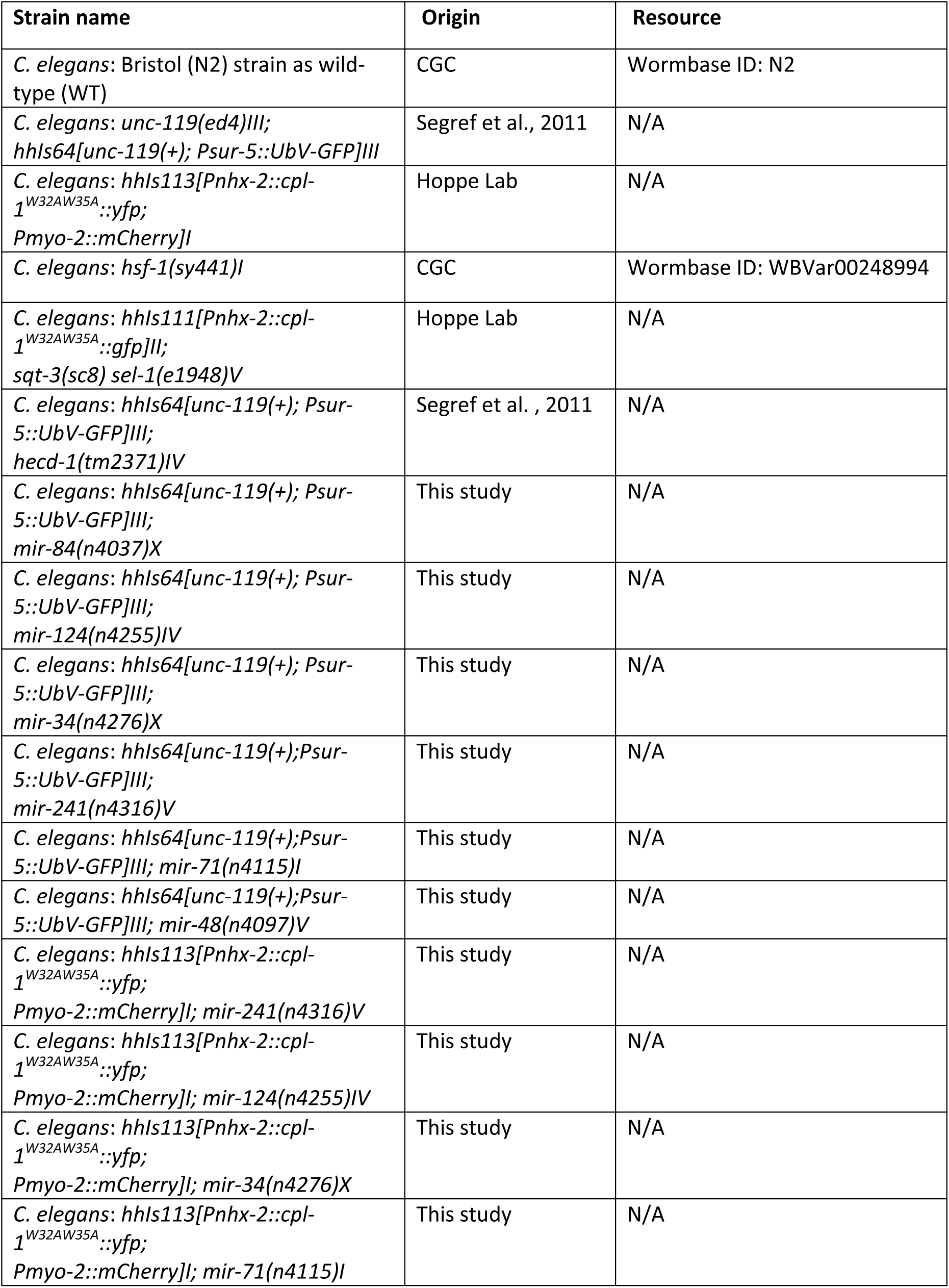
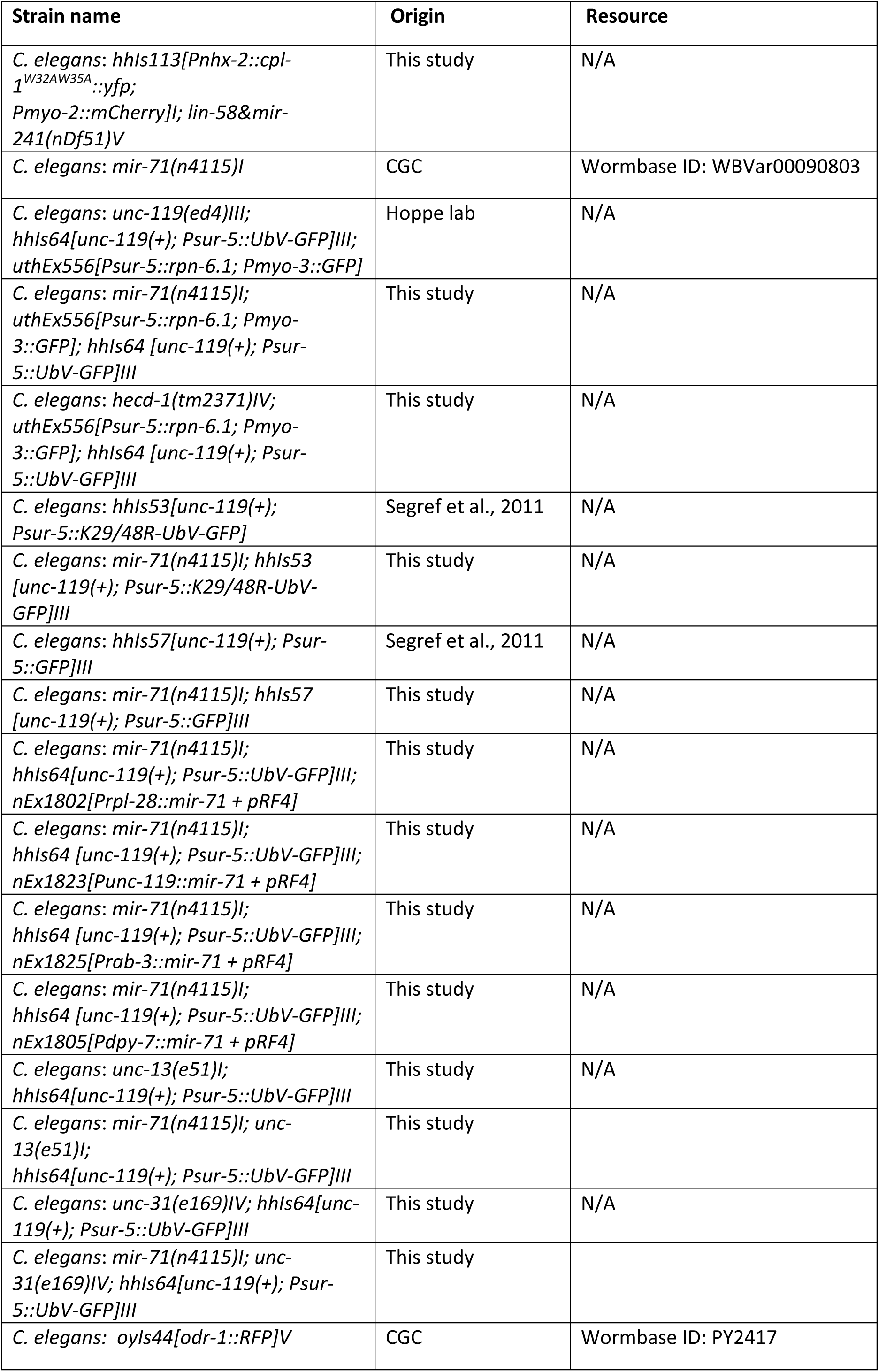
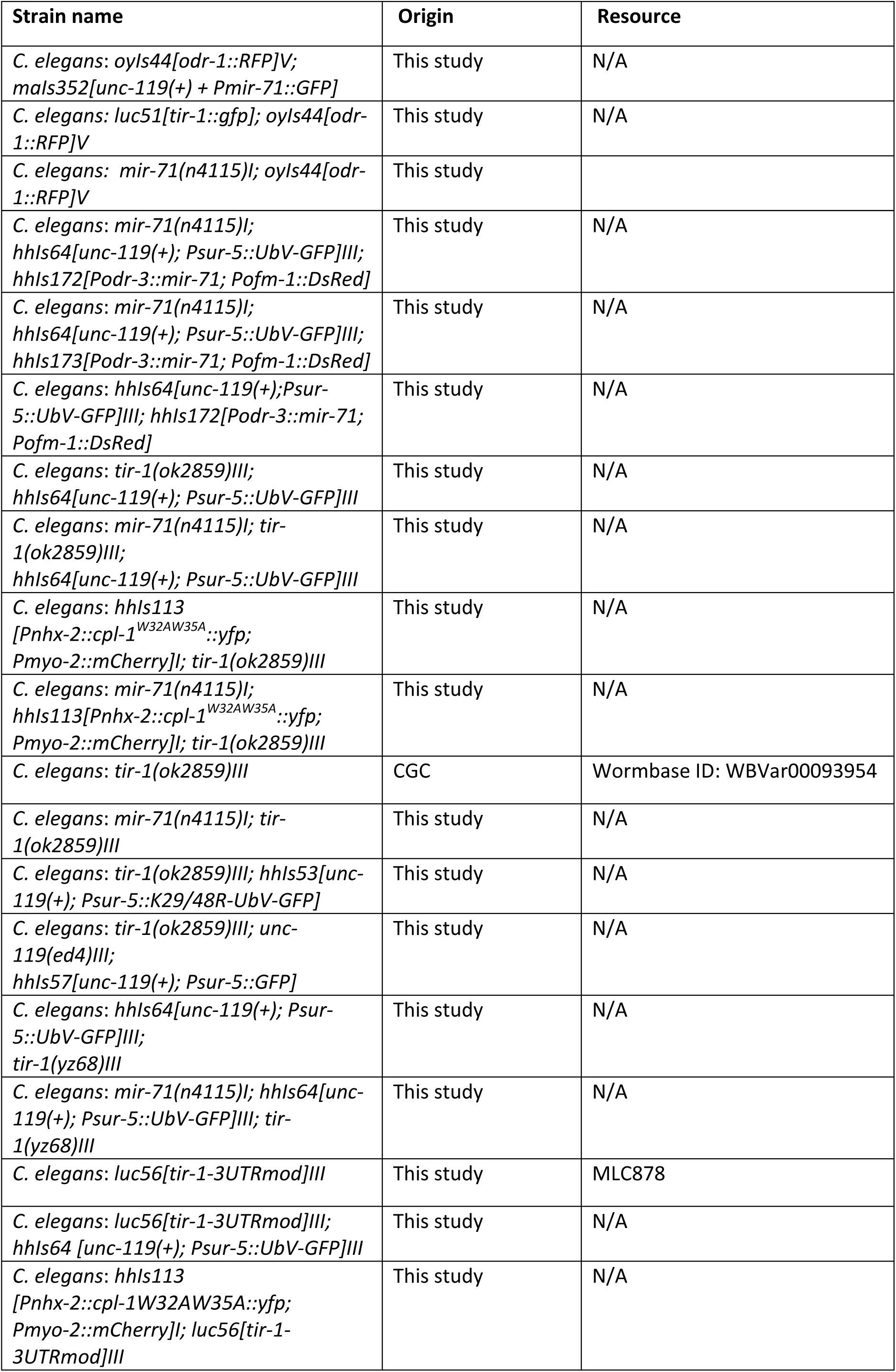
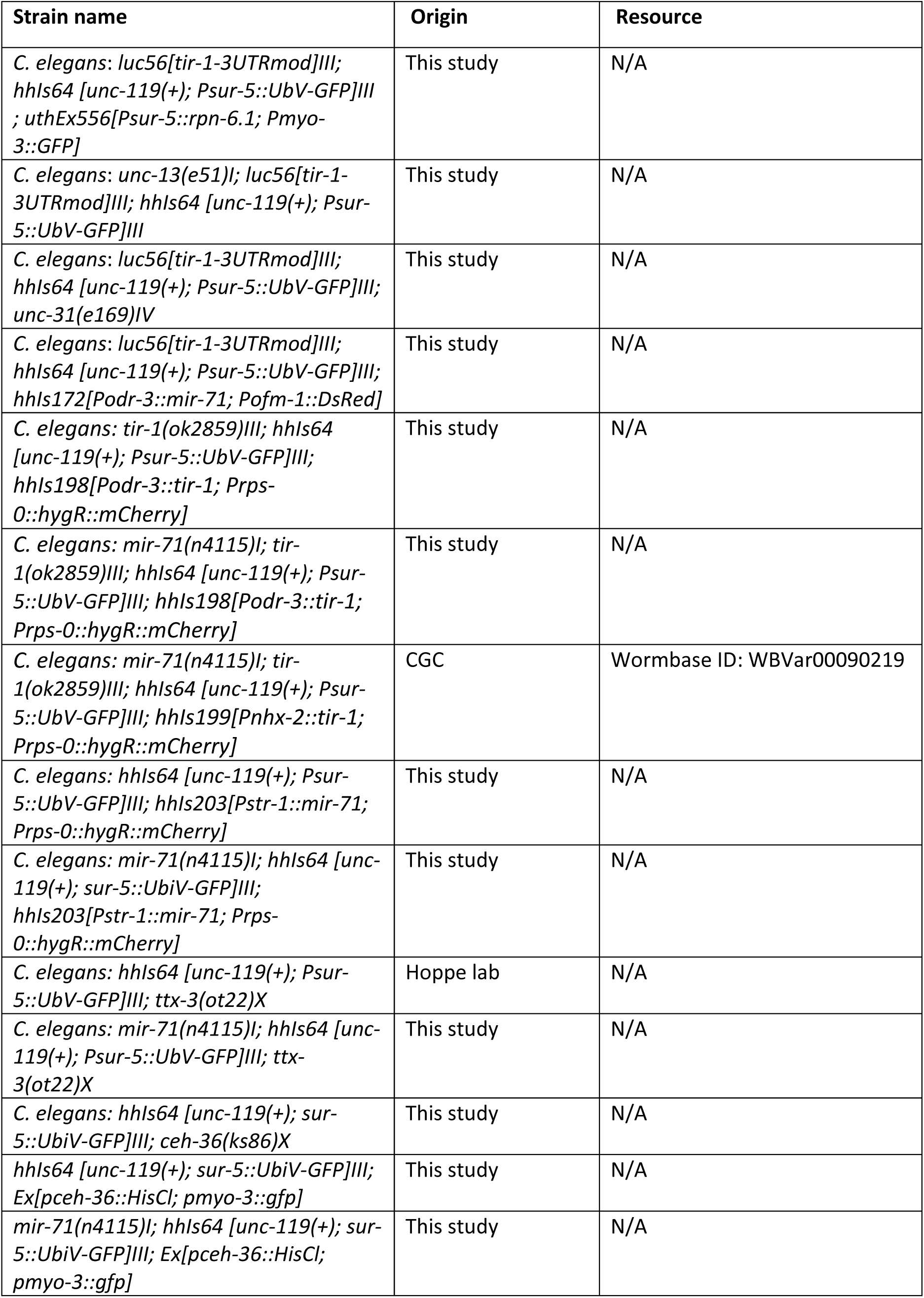
| *C. elegans* strains used in this study.

**Supplementary Table 7.**
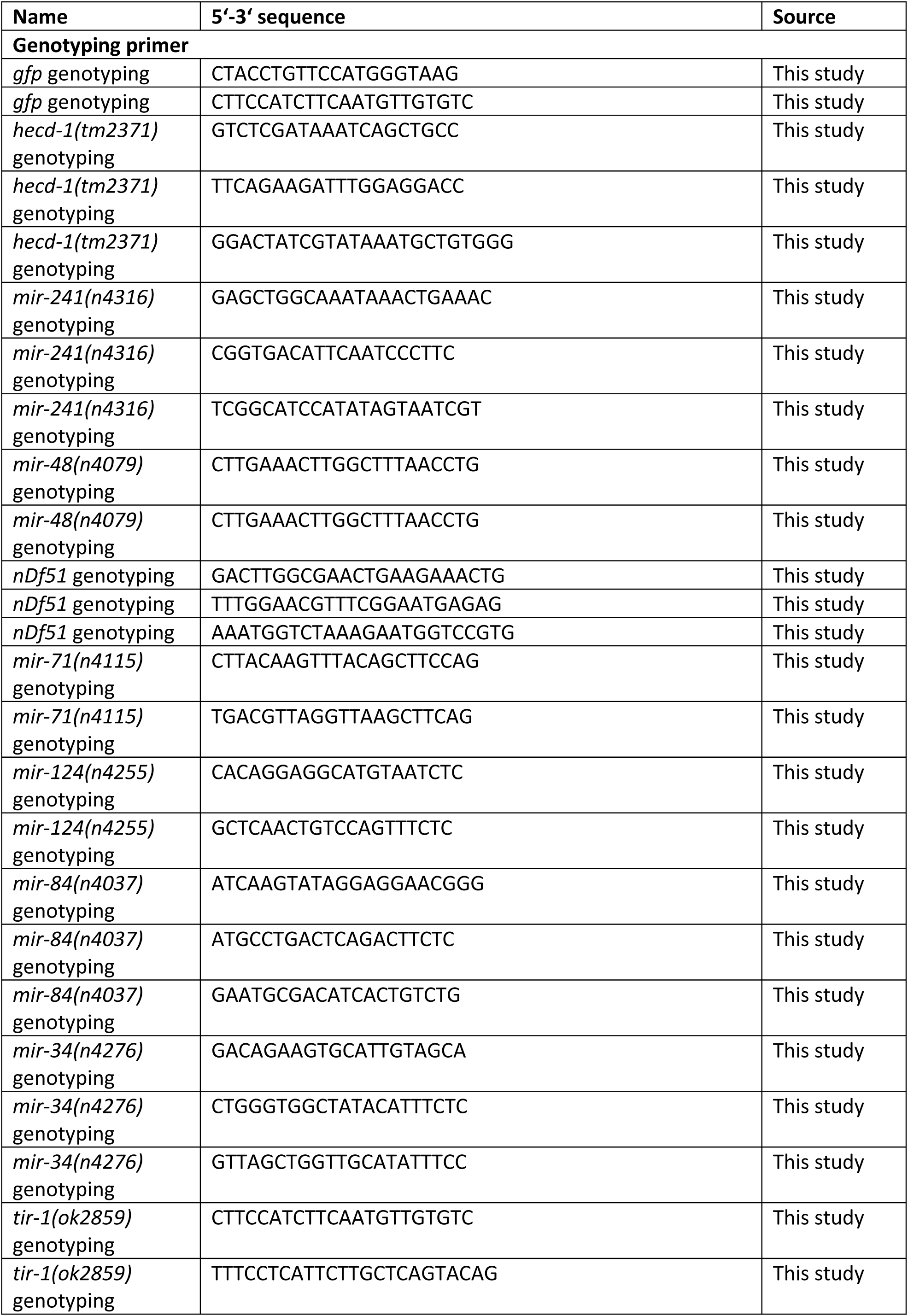
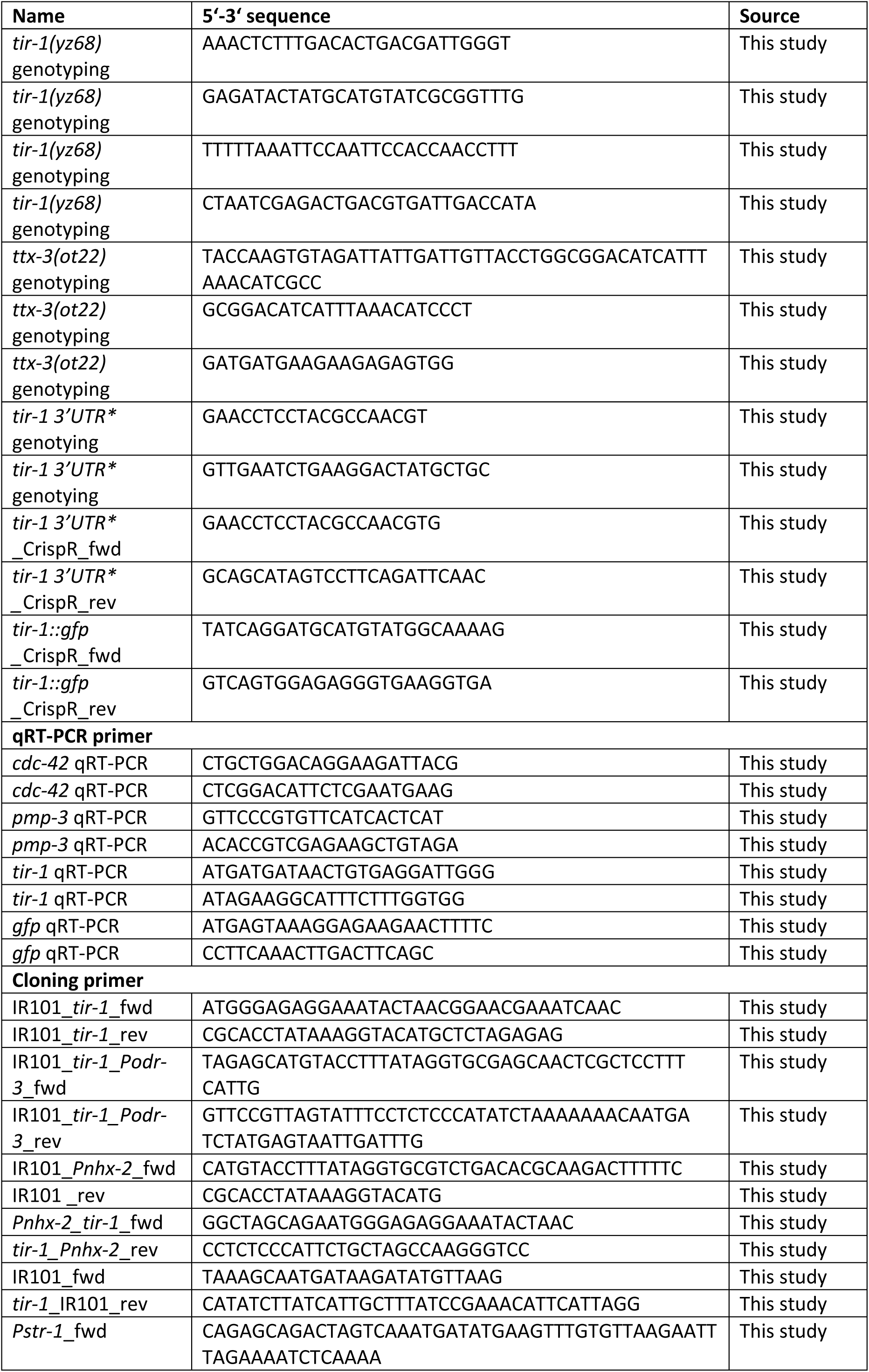
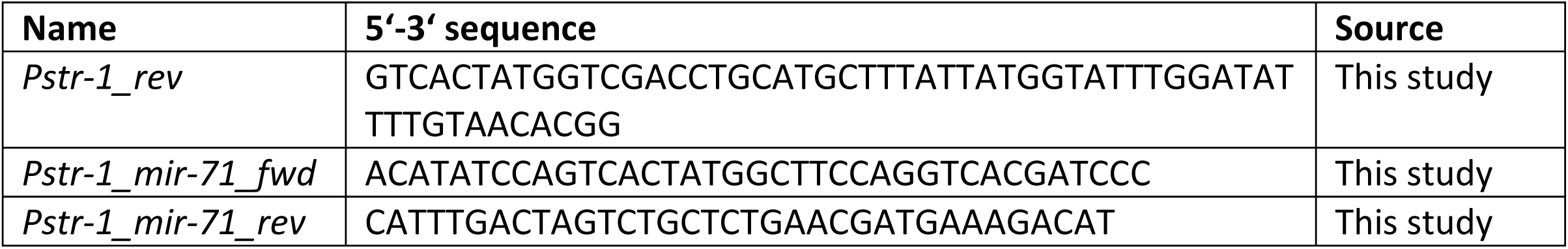
| Oligonucleotides used in this study.

**Supplementary Table 8.**
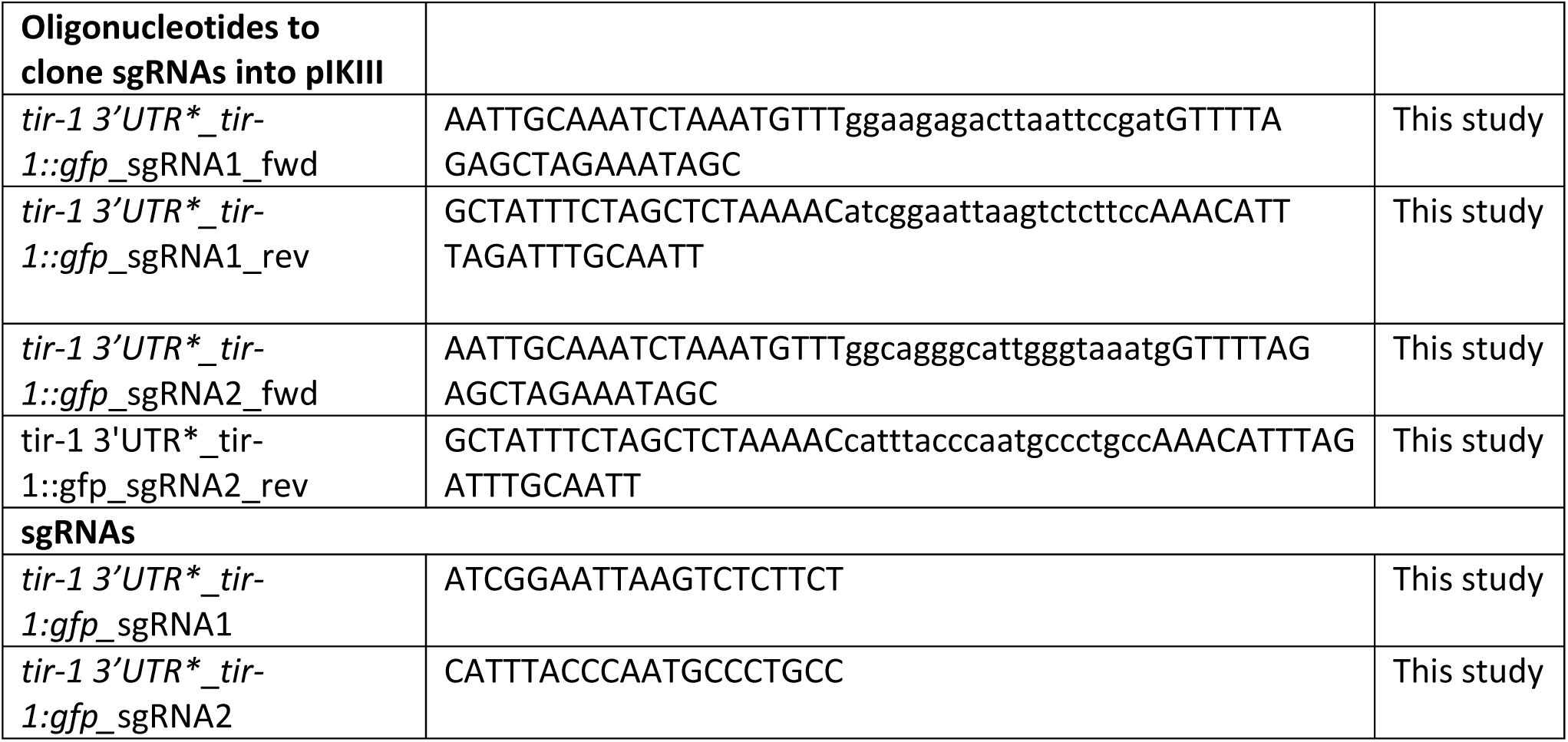
| Oligonucleotides and repair templates used for CrispR/Cas9-mediated gene editing.

## CrispR/Cas9 repair templates

### *tir-1 3’UTR* mutant

TTCACAAAATATTATATTGCACGAGCACAAAAATTTGAGAATACGTAATGGTAACAAAGAAAACTACAGTACCTCCTTTAGTGACTACTATAGCTCTTTATGTGCTCGATTTTCGAAGGTTTTATCAATTATATCTTTTATTTTCAGATGCCTTCTATCTCCCGAAAAACTACCCAACAACGGTGGCAGACCACAAATACAGTAAGCCGTACTGGGCCATCCAGAAGCATCGGTGGACCAAGAATGGAACCTCCTACGCCAACGTGTAAGTTGTTGAAACATCGTCACTTGGTATATTATTTAATTTTTTTAGTCTTTTCAGTTACACCAACTGGATCACAAGAACGTGCAACATCGACGAGAAGGAAAATTCAGCCTTCTGCATCCACAACTTCTGATCGGAATTAAGTCTCTTCTTTAATCAATACATCAAGTGCCGGTTCATTTTATCCATTTTCCAACAAAAAACCAATTTTCTTCAACCATCCCAATATCAAATATCAAACTCCCAATGGTACCATCATCCCACACGAAAGTTTTAATGTATTTGACCCTCATTTTAAATCAGTCAAAACCTCAAAAAAAGTGTTCCACAAAAAACTGGAAATTCAAAGCGTAGAATCTATGATTTCAACTGTCTCTCTCTCTTCGTCATTTACCCAATGCCCTGCCGTTTTCTGCAAATAATAATAATAATTGTAATAATCTCTATAAATTATGTGATTGATATGAAACCAAAAAGAAAACTCTGAAACTTCACTTTCTTATAAGGTACCTGTTTATATATTCCGATCTCCATCTGTTTCATTCGTTTCACTTTTTCTGTTTTATTTCGTGCAACGTGTAATAGTCGATCAACGCCCAACTCAGGAACCCATCAAATTCGAACCCTTTAACTCCCTCTTTAAATTGAACTTTTTATCTTAACAATTACCGGGTCTAACACCCTTCCAGATTTCTAATGTTATACCTAATGAATGTTTCGGATTATTGGGGATCCTCTAGAGTCGACCTGCAGGCATGCAAGCTTGGCGTAATCATGGTCATAGCTGTTTCCTGTGTGAAATTGTTATCCGCTCACAATTCCACACAACATACGAGCCGGAAGCATAAAGTGTAAAG CCTGGGGTGCCTAATGAGTGAGCTAACTCACATTAATTGCGTTGCGCTCACTGCCCGCTTTCCAGTCGGGAAACCTGTCGTGCCAGCTGCATTAATGAATCGGCCAACGCGCGGGGAGAGGCGGTTTGCGTATTGGGCGCTCTTCCGCTTCCTCGCTCACTGACTCGCTGCGCTCGGTCGTTCGGCTGCGGCGAGCGGTATCAGCTCACTCAAAGGCGGTAATACGGTTATCCACAGAATCAGGGGATAACGCAGGAAAGAACATGTGAGCAAAAGGCCAGCAAAAGGCCAGGAACCGTAAAAAGGCCGCGTTGCTGGCGTTTTTCCATAGGCTCCGCCCCCCTGACGAGCATCACAAAAATCGACGCTCAAGTCAGAGGTGGCGAAACCCGACAGGACTATAAAGATACCAGGCGTTTCCCCCTGGAAGCTCCCTCGTGCGCTCTCCTGTTCCGACCCTGCCGCTTACCGGATACCTGTCCGCCTTTCTCCCTTCGGGAAGCGTGGCGCTTTCTCATAGCTCACGCTGTAGGTATCTCAGTTCGGTGTAGGTCGTTCGCTCCAAGCTGGGCTGTGTGCACGAACCCCCCGTTCAGCCCGACCGCTGCGCCTTATCCGGTAACTATCGTCTTGAGTCCAACCCGGTAAGACACGACTTATCGCCACTGGCAGCAGCCACTGGTAACAGGATTAGCAGAGCGAGGTATGTAGGCGGTGCTACAGAGTTCTTGAAGTGGTGGCCTAACTACGGCTACACTAGAAGAACAGTATTTGGTATCTGCGCTCTGCTGAAGCCAGTTACCTTCGGAAAAAGAGTTGGTAGCTCTTGATCCGGCAAACAAACCACCGCTGGTAGCGGTGGTTTTTTTGTTTGCAAGCAGCAGATTACGCGCAGAAAAAAAGGATCTCAAGAAGATCCTTTGATCTTTTCTACGGGGTCTGACGCTCAGTGGAACGAAAACTCACGTTAAGGGATTTTGGTCATGAGATTATCAAAAAGGATCTTCACCTAGATCCTTTTAAATTAAAAATGAAGTTTTAAATCAATCTAAAGTATATATGAGTAAACTTGGTCTGACAGTTACCAATGCTTAATCAGTGAGGCACCTATCTCAGCGATCTGTCTATTTCGTTCATCCATAGTTGCCTGACTCCCCGTCGTGTAGATAACTACGATACGGGAGGGCTTACCATCTGGCCCCAGTGCTGCAATGATACCGCGAGACCCACGCTCACCGGCTCCAGATTTATCAGCAATAAACCAGCCAGCCGGAAGGGCCGAGCGCAGAAGTGGTCCTGCAACTTTATCCGCCTCCATCCAGTCTATTAATTGTTGCCGGGAAGCTAGAGTAAGTAGTTCGCCAGTTAATAGTTTGCGCAACGTTGTTGCCATTGCTACAGGCATCGTGGTGTCACGCTCGTCGTTTGGTATGGCTTCATTCAGCTCCGGTTCCCAACGATCAAGGCGAGTTACATGATCCCCCATGTTGTGCAAAAAAGCGGTTAGCTCCTTCGGTCCTCCGATCGTTGTCAGAAGTAAGTTGGCCGCAGTGTTATCACTCATGGTTATGGCAGCACTGCATAATTCTCTTACTGTCATGCCATCCGTAAGATGCTTTTCTGTGACTGGTGAGTACTCAACCAAGTCATTCTGAGAATAGTGTATGCGGCGACCGAGTTGCTCTTGCCCGGCGTCAATACGGGATAATACCGCGCCACATAGCAGAACTTTAAAAGTGCTCATCATTGGAAAACGTTCTTCGGGGCGAAAACTCTCAAGGATCTTACCGCTGTTGAGATCCAGTTCGATGTAACCCACTCGTGCACCCAACTGATCTTCAGCATCTTTTACTTTCACCAGCGTTTCTGGGTGAGCAAAAACAGGAAGGCAAAATGCCGCAAAAAAGGGAATAAGGGCGACACGGAAATGTTGAATACTCATACTCTTCCTTTTTCAATATTATTGAAGCATTTATCAGGGTTATTGTCTCATGAGCGGATACATATTTGAATGTATTTAGAAAAATAAACAAATAGGGGTTCCGCGCACATTTCCCCGAAAAGTGCCACCTGACGTCTAAGAAACCATTATTATCATGACATTAACCTATAAAAATAGGCGTATCACGAGGCCCTTTCGTC

### tir-1∷gfp

TCACAAAATATTATATTGCACGAGCACAAAAATTTGAGAATACGTAATGGTAACAAAGAAAACTACAGTACCTCCTTTAGTGACTACTATAGCTCTTTATGTGCTCGATTTTCGAAGGTTTTATCAATTATATCTTTTATTTTCAGATGCCTTCTATCTCCCGAAAAACTACCCAACAACGGTGGCAGACCACAAATACAGTAAGCCGTACTGGGCCATCCAGAAGCATCGGTGGACCAAGAATGGAACCTCCTACGCCAACGTGTAAGTTGTTGAAACATCGTCACTTGGTATATTATTTAATTTTTTTAGTCTTTTCAGTTACACCAACTGGATCACAAGAACGTGCAACATCGACGAGAAGGAAAATTCAGCCTTCTGCATCCACAACTTCTGATCGGAATATGAGTAAAGGAGAAGAACTTTTCACTGGAGTTGTCCCAATTCTTGTTGAATTAGATGGTGATGTTAATGGGCACAAATTTTCTGTCAGTGGAGAGGGTGAAGGTGATGCAACATACGGAAAACTTACCCTTAAATTTATTTGCACTACTGGAAAACTACCTGTTCCATGGGTAAGTTTAAACATATATATACTAACTAACCCTGATTATTTAAATTTTCAGCCAACACTTGTCACTACTTTCTGTTATGGTGTTCAATGCTTCTCGAGATACCCAGATCATATGAAACGGCATGACTTTTTCAAGAGTGCCATGCCCGAAGGTTATGTACAGGAAAGAACTATATTTTTCAAAGATGACGGGAACTACAAGACACGTAAGTTTAAACAGTTCGGTACTAACTAACCATACATATTTAAATTTTCAGGTGCTGAAGTCAAGTTTGAAGGTGATACCCTTGTTAATAGAATCGAGTTAAAAGGTATTGATTTTAAAGAAGATGGAAACATTCTTGGACACAAATTGGAATACAACTATAACTCACACAATGTATACATCATGGCAGACAAACAAAAGAATGGAATCAAAGTTGTAAGTTTAAACATGATTTTACTAACTAACTAATCTGATTTAAATTTTCAGAACTTCAAAATTAGACACAACATTGAAGATGGAAGCGTTCAACTAGCAGACCATTATCAACAAAATACTCCAATTGGCGATGGCCCTGTCCTTTTACCAGACAACCATTACCTGTCCACACAATCTGCCCTTTCGAAAGATCCCAACGAAAAGAGAGACCACATGGTCCTTCTTGAGTTTGTAACAGCTGCTGGGATTACACATGGCATGGATGAACTATACAAATAGGTCTCTTCTTTAATCAATACATCAAGTGCCGGTTCATTTTATCCATTTTCCAACAAAAAACCAATTTTCTTCAACCATCCCAATATCAAATATCAAACTCCCAATGTCTTTCAATCATCCCACACGAAAGTTTTAATGTATTTGACCCTCATTTTAAATCAGTCAAAACCTCAAAAAAAGTGTTCCACAAAAAACTGGAAATTCAAAGCGTAGAATCTATGATTTCAACTGTCTCTCTCTCTTCCTCATTTACCCAATGCCCTGCCGTTTTCTGCAAATAATAATAATAATTGTAATAATCTCTATAAATTATGTGATTGATGTCTTTATGAAACCAAAAAGAAAACTCTGAAACTTCACTTTCTTATAATCTTTCTGTTTATATATTCCGATCTCCATCTGTTTCATTCGTTTCACTTTTTCTGTTTTATTTCCTGCAACGTGTAATAGTCGATCAACCCCCAACTCAGGAACCCATCAAATTCGAACCCTTTAACTCCCTCTTTAAATTGAACTTTTTATCTTAACAATTACCGGGTCTAACACCCTTCCAGATTTCTAATGTTATACCTAATGAATGTTTCGGATTA TT

